# FastACI: a Toolbox for Investigating Auditory Perception using Reverse Correlation

**DOI:** 10.1101/2025.06.01.657236

**Authors:** Alejandro Osses, Azal Le Bagousse, Léo Varnet

## Abstract

The fastACI toolbox provides a compilation of tools for collecting and analyzing data from auditory reverse-correlation experiments. These experiments involve behavioral listening tasks including one or more target sounds presented with some random fluctuation, typically in the form of additive background noise. In turn, the paired stimulus-response data from each trial can be used to assess the relevant acoustic features that were effectively used by the listener while performing the task. The results are summarized as a matrix of perceptual weights termed auditory classification image. The framework provided by the toolbox is flexible and it has been so far used to probe different auditory mechanisms such as tone-in-noise detection, amplitude modulation detection, phoneme-in-noise categorization, and word segmentation. In this article, we present the structure of the toolbox, how it can be used to run existing experiments or design new ones, as well as the main options for analyzing the collected data. We then illustrate the capabilities of the toolbox through five case studies: a replication of a pioneering reverse correlation study from 1975, an example of reproduction of the analyses of one of our previous studies, a comparison of the results of three phoneme-categorization experiments, and a quantification of how noise type and estimation method affect the quality of the resulting auditory classification image.

## 1 INTRODUCTION

Auditory reverse correlation (revcorr) is a psychophysical paradigm that allows to determine which acoustic features in the test stimuli are effectively used as cues by participants during a listening experiment, with only minimal prior assumptions. This method relies on two critical ingredients: (1) the introduction of random fluctuations into the stimulus (such as background noise) and (2) the trial-by-trial (“molecular,” or “microscopic”) analysis of the relationship between the specific noise samples and the corresponding participant responses (Neri, 2018; Murray, 2011). By examining how specific noise fluctuations drive specific responses from the listener or “observer,” this technique provides a valuable insight into the perceptual process, that is not accessible through classic (“macroscopic,” sometimes also referred to as “molar”) psychophysics based on averaging over hundreds of trials.

The concept of a molecular approach was initially theorized by David Green, stating that “The development of some form of molecular psychophysics seems as inevitable as the development of more quantitative theories of sensory functions. Indeed, more and more crucial tests of such theories will be possible on the molecular level as they become more exact and quantitative” (Green, 1964). Less than a decade later, this prediction was proved to be correct when Ahumada and Lovell applied the revcorr paradigm for the first time in a series of two experiments focusing on the ability to detect a pure tone in a white Gaussian noise masker. They applied a multiple regression analysis to the spectral (Ahumada and Lovell, 1971) or spectrotemporal (Ahumada et al., 1975) representation of the noise in each trial to estimate the contribution of different auditory features to the listeners’ decision regarding the presence or absence of the tone. Their results showed that, in a tone-in-noise detection task, the greatest perceptual weight is assigned to the signal frequency, with negative weights at frequencies above and below the signal frequency, and immediately before the signal. We present a replication of these results in Section 6.1.

Since Ahumada’s seminal studies, the reverse correlation approach became very popular in psychoacoustic research. Recent applications include studies on loudness perception (Ponsot et al., 2013), modulation perception (Varnet and Lorenzi, 2022; Ponsot et al., 2021), phoneme-in-noise perception (Varnet et al., 2013, 2015a; Osses and Varnet, 2024), word segmentation (Osses et al., 2023), and perception of paralinguistic prosodic features (Ponsot et al., 2018; Goupil et al., 2021). The results of reverse correlation experiments are typically displayed as matrices of time-frequency weights, sometimes referred to as auditory classification images (ACIs). For this reason, we will use the terms “reverse correlation method,” “revcorr method,” and “ACI method” interchangeably throughout this text.^1^

The core principle of the ACI method is to correlate observer decisions with noisy stimulus features over large sets of stimuli. Beyond this, the methodological details are left to the experimenter’s discretion. For instance, the task could involve detection or discrimination; noise levels could be fixed or adaptive; and there could be one or multiple targets. Similarly, several methods have been proposed for estimating the perceptual weights, including correlation, logistic regression, or penalized regression. All these specific experimental designs and analysis schemes can be incorporated into a revcor experiment, provided that the noise waveforms presented in each trial are recorded along with the corresponding participant responses.

In this article, we introduce a framework for conducting listening experiments and post-processing experimental data using the reverse correlation method. Our primary motivation to develop a new toolbox originated from the need to store all individual waveforms used during the experimental sessions to derive the ACI weights. Other well-established psychophysics tools, such as those provided in the AFC (Ewert, 2013) or APEX toolboxes (Francart et al., 2008) require the specification of target sounds, but typically generate background noises on the fly, without tracking the specific waveform presented with the target stimuli. Another approach is adopted in tools such as the CLEESE toolbox (Burred et al., 2019). However, while CLEESE provides a convenient means to generate and store speech stimuli with random fluctuations in prosody, it does not provide specific tools for running the experiment or analyzing the collected data. A secondary motivation for creating a new toolbox was therefore to integrate data analysis tools within the same framework as the experiment. During data post-processing, the revcorr method involves reading the labeled responses and linking them with the dimensions of the test stimuli representations, which are usually time and frequency. We therefore decided to compile the required tools within a single framework, enabling transparent replication of previous studies and reproducibility of analyses.

We aimed to make this toolbox a turnkey solution for conducting revcorr experiments: installing the toolbox, setting up, running, and analyzing a simple experiment should require minimal effort from the experimenter. Another central objective was to keep the framework flexible, allowing a straightforward extensibility in future research. Historically, the revcorr method has required many trial presentations— often in the order of thousands—to derive clear time-frequency ACI weights. One long-term goal of the project is to gradually reduce the number of trials required to obtain ACIs. For this reason, we decided to name the toolbox “fastACI”. More generally, the fastACI toolbox can serve for multiple purposes, allowing users to:

1. Conduct listening experiments, based on a single-interval yes/no task or a two-interval forced choice, with one independent variable (e.g., signal-to-noise ratio) that can be either adjusted using an adaptive procedure or held constant at a predetermined value using a constant-stimulus procedure.
2. Run listening experiments involving either human or artificial listeners.
3. Automatically regenerate target sounds and background noises corresponding to an experiment, even if the local waveforms are no longer available.
4. Post-process the data collected on a listener using several available statistical models to derive the time-frequency ACI weights for this listener.
5. Provide an open-source framework for facilitating replication and computational reproducibility in the field of auditory revcorr.

In the following sections, we describe the general structure of the fastACI toolbox (Section 2), how to run an experiment or design new ones (Section 3), the conventions used for storing the data (Section 4), and how to post-process the collected data (Section 5). In a final section (Section 6) we illustrate the possibilities of the toolbox through five case studies.

## 2 STRUCTURE OF THE TOOLBOX

The fastACI toolbox is a command-line based set of tools coded in Matlab and openly available on Zenodo (Osses and Varnet, 2021b) and hosted on GitHub (https://github.com/LeoVarnet/fastACI). It has been tested with versions between R2012b and R2024a on Windows, Linux, and Mac. It was developed as an extension of the ACI toolbox, published in 2015 (Varnet, 2015). In its new version, it borrows coding conventions from two other packages, the AMT toolbox (Majdak et al., 2022) for the management of parameters and naming of functions and the AFC toolbox (Ewert, 2013) for the definition of experiments.

The fastACI toolbox—hereafter referred to as “the toolbox”—is based on two main modules for data collection and data post-processing using the reverse-correlation method. The main functions for these two modules are `fastACI_experiment.m` and `fastACI_getACI.m`, respectively, as indicated in the block diagram of Fig. 1.

**Figure 1.**
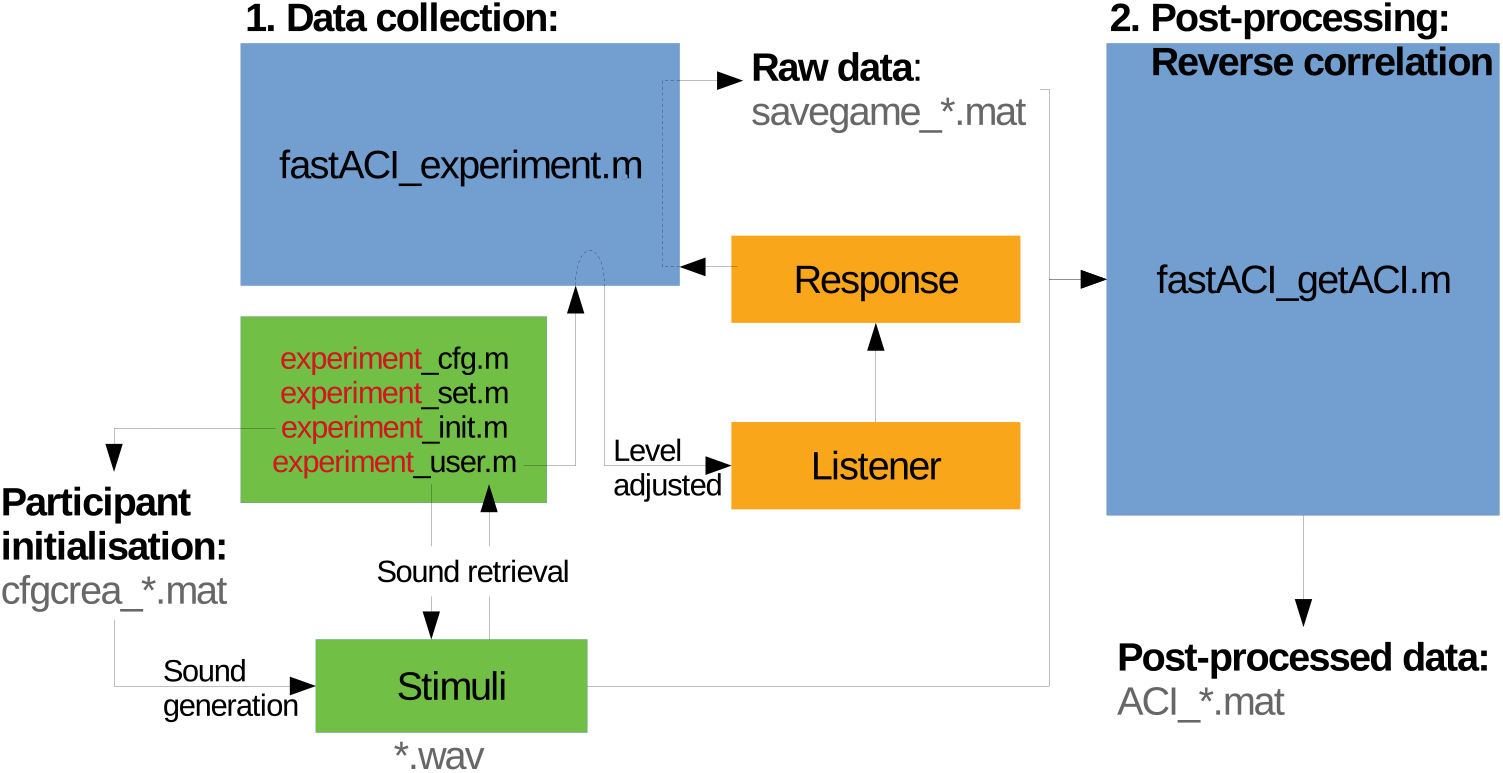
Block diagram of the data flow in the fastACI toolbox for an experiment called ‘experiment.’ The blue blocks represent base functions and subfunctions from the fastACI environment. The green blocks represent the experiment definition, consisting of four scripts (`*_cfg.m`, `*_set.m`, `*_init.m`, `_user.m`. The orange blocks represent the participants to the experiment. The binary data generated during the experimental data collection or post-processing are indicated in grey text. More details are given in the text.

The data collection module, controlled by the function `fastACI_experiment.m`, manages the entire workflow of an auditory experiment, from generating the auditory stimuli to presenting them to participants and recording the behavioral responses (see Section 3). This module relies on four experiment-specific configuration scripts—namely, `expName_cfg.m`, `expName set.m`, `expName_init.m`, and `expName_user.m`— which specify the parameters of the experimental protocol for the experiment with the custom name “expName.” This provides an easy way for the user to customize the experiment, such as selecting different target sounds, turning off the feedback or the training (warm-up) session, or modifying the rules of the adaptive procedure. The stimuli are first generated and stored in a participant-specific folder, along with a .mat file *cfgcrea *.mat* summarizing all parameters used during stimulus generation. During the data collection phase, the auditory stimuli are retrieved from this directory and played back to the participants. The system adjusts the sound level as well as any other specified parameter—typically, the signal-to-noise ratio (SNR)—to meet the experimental requirements. Participant responses are recorded and saved in a .mat files (*savegame *.mat*), along with all parameters relevant to the experimental session, ensuring reproducibility and transparency in data collection (see Section 4).

The data post-processing module of the toolbox is managed by the function `fastACI_getACI.m`, which implements the revecorr analysis of the collected data (see Section 5). The function takes as input the raw response data stored in the *savegame *.mat* file, along with several optional parameters that specify the details of the analysis. The stimuli are either retrieved from the participant folder or re-generated on the fly, and are then converted into a particular matrix representation. This matrix representation is typically a spectro-temporal representation, that can be changed by defining an experiment-specific `expName_dataload.m` function. These representations are then analyzed together with the corresponding participant’s responses. The outcome of this analysis is stored as a post-processed data file (*ACI_*.mat*). The modular design allows researchers to apply different analysis techniques or modify parameters within `fastACI_getACI.m` to explore various aspects of the results, making it a versatile tool for auditory research.

## 3 RUNNING AN EXPERIMENT

### 3.1 First-time use

Once downloaded from Zenodo (Osses and Varnet, 2021b), the fastACI toolbox can be initialized by running `startup_fastACI.m`. This script automatically adds all the necessary directories to the Matlab path for the duration of the current session, and checks for the required data folders (*dir data* and *dir datapost*) and dependencies. Two third-party toolboxes are mandatory: the AMT toolbox (Majdak et al., 2022) and the LTFAT toolbox (Søndergaard et al., 2012) (included within AMT). Additionally, several optional toolboxes may be used, such as the AFC toolbox (Ewert, 2013), the PhaseRet toolbox (Průša, 2017) for generating MPS noise or bump noise (see Section 3.3), Praat (Boersma and Weenink, 2025) for analyzing the spectral content of speech stimuli, and WORLD (Morise, 2016; Morise et al., 2016) for dimensional noise approaches (not described here, see Osses et al., 2023, for more details).

### 3.2 Running a pre-existing experiment

The toolbox offers a range of predefined experiments available natively. The scripts describing each of these experiments are stored in a separate folder under the directory .*/fastACI-main/Experiments/*. A list of predefined experiments is provided in Table 1.

**Table 1.** List of published fastACI experiments, sorted by date of publication (more recent first). AM = Amplitude modulation.

| Experiment name | Task | Target sounds | Reference |
| --- | --- | --- | --- |
| replication_ahumada1975 | tone-in-noise detection | 100-ms 500-Hz pure tone or silence | Le Bagousse and Varnet (2025)<br>Ahumada et al. (1975) |
| speechACI_Logatome | phoneme categorization | Pairs of phonetic contrasts using /aba/, /ada/, /aga/, /apa/, /ata/ from the same male speaker | Osses and Varnet (2024)<br>Carranante et al. (2024) |
| segmentation | word segmentation | Pairs of homophonic sentences in French (e.g., “c’est l’amie” / “c’est la mie”) | Osses et al. (2023) |
| modulationACI | AM detection | 4-Hz modulated tone or pure tone (1 kHz carrier) | Varnet and Lorenzi (2022) |
| speechACI_varnet2013 | phoneme categorization | /aba/-/ada/, female speaker | Osses and Varnet (2021a)<br>Varnet et al. (2013) |
| speechACI_varnet2015 | phoneme categorization | /alda/-/alga/-/arda/-/arga/, male speaker | Varnet et al. (2016)<br>Varnet et al. (2015a)<br>Varnet et al. (2015b) |

An experiment can be run using function `fastACI_experiment.m`, which requires as input arguments the participant ID, the experiment name, and, optionally, the condition to be tested. For instance, in order to start experiment speechACI_varnet2013 for participant ‘S01’ using a white noise masker, the appropriate command is: `fastACI_experiment(`‘`speechACI_varnet2013`’,’`S01`’,’`white`’).

When running `fastACI_experiment.m`, it is first checked whether the participant is being run for the first time (function `Check_local_dir_data.m`). If previous sessions are found, then the next trial is resumed assuming that all stimuli are already on disk. If no previous session is found, the participant is first initialized (function `fastACI_experiment_init.m`) before the first session can start. In particular, all target and masker waveforms are generated and stored in a participant-specific directory, together with a *cfgcrea*_*\*.mat* file containing all configuration settings for the experiment being ran (see Section 4). Because of the large number of waveforms required for running the experiments, this file also stores the seed numbers used to generate all the noise waveforms. This feature enables the toolbox to retrieve the exact same noise waveforms at any moment if the local stimuli are removed from the computer. This action is automatically performed if a previously created *cfgcrea*_*\*.mat* file is found on disk without finding the associated waveforms (see Section 4.4).

Once the experiment is initialized (or resumed), the script `fastACI_experiment.m` takes care of running the test, collecting and storing the participant’s data, and adjusting the experimental variable (expvar) from one trial to the next. The function `fastACI_trial_current.m`, that describes the structure of a trial, is iteratively called until the last trial of the session or of the experiment is reached, or the participant requires a break.

The function `fastACI_trial_current.m` is central to the toolbox. It takes as input the parameters of the experiment (stored in the *cfg_game.mat*) and current state of the experiment (the structure data_passation), executes a single stimulus-response trial, and updates the data_passation structure. During the trial, it displays relevant information on-screen for the participant such as the trial number, upcoming breaks, and available response options. The stimulus—or stimuli in the case of a two-interval task— is generated using the experiment-specific `*_user.m` function (see Section 3.3) and played back using the `audioplayer.m` function from Matlab. The function `Response_keyboard.m` then displays the different response alternatives on screen and waits for an input of the participant. There are typically three possible answers: the names of the two target sounds (by default the names of the wavefiles, but we encourage experimenters to overwrite this default using the ‘response names’ field of the cfg_crea structure) and ‘press 3 to take a break’. In case of a two-interval forced-choice task, the first two response options are ‘X first and Y second’ and ‘Y first and X second’, with X and Y the names of the two target sounds. The participant’s response is stored in the data_passation structure together with the target actually presented, the current value of expvar and the response time. Finally, the value of expvar is updated, if needed, according to the specified staircase rules. The information displayed on screen (e.g., feedback, instructions) is largely customizable, and the most difficult experiment may also include probe stimuli (easy stimuli presented periodically after *N* trials) or a training (warm-up) session. If a training session is requested, the `Response_keyboard.m` function displays four additional options: listening to the original noise-free targets, listening to the noisy stimulus again, or leaving the warm-up session to start the main experiment. During the warm-up session, a feedback on the answer is automatically provided before the next trial begins.

### 3.3 Running a new experiment

As indicated by the green blocks in Fig. 1, an experiment is implemented by defining a compulsory number of four scripts that are named with the experiment name (“expName”) as prefix. For instance, for experiment speechACI Logatome, these scripts are:

- `speechACI_Logatome_cfg.m`,
- `speechACI_Logatome_user.m`,
- `speechACI_Logatome_set.m`,
- `speechACI_Logatome_init.m`.

In general, we will refer to these scripts as the configuration (`*_cfg.m`), user (`*_user.m`), set-up (`*_set.m`) and initialization (`*_init.m`) files. This experiment structure was inspired by the definitions in the AFC toolbox (Ewert, 2013). The experiment files are briefly explained in order of execution below:

* _**set.m**: The set-up file contains the definition of variables that do not change during the experiment. There are no compulsory variables to be defined here, but we recommend specifying variables such as sampling frequency (cfg_game.fs), presentation level (cfg_game.SPL), calibration level of the waveforms (cfg_game.dBFS) and number of targets stimuli (cfg_game.N_target). In this script, we also provide the possibility to overwrite default parameters, such as the calibration level of the playback (by default equal to cfg_game.dBFS) or the number of total trials (cfg_game.N).
* _**init.m**: The initialization file generates the *cfgcrea*_*\*.mat* file and, if the sound stimuli are not yet stored on disk, prepares the target sounds and generates the background noises. This script is only run once, at the beginning of the experimental data collection. The default method for generating noise waveforms is through the `Generate_noise.m` function, although this can be customized as needed in the initialization file. Several predefined noise types are available, including white noise (‘white’), pink noise (‘pink’), bump noise (‘bumpv1p2 10dB’), and MPS noise (‘sMPSv1p3’). The last two options correspond to maskers with a flat long-term averaged spectrum, similar to white noise, but exhibiting larger random envelope fluctuations (see Osses and Varnet, 2024, for a more detailed description). As discussed in Section 6.4, the enhanced fluctuations present in bump and MPS noise make them more efficient than white noise for deriving an ACI.
* _**cfg.m**: The configuration file contains all remaining details for setting up the experiment. It is executed once at the beginning of each experiment. This file defines the entire experimental configuration, including mandatory parameters such as the experimental variable expvar, whether it should be changed adaptively from one trial to the next (cfg_game.adapt), its initial value at the beginning of each experimental session (cfg_game.startvar), and the number of trials per session (cfg_game.sessionN). In most cases, the experimental variable corresponds to the stimulus dimension that is systematically varied during the experiment. For example, in speech-in-noise tests, the stimuli are adjusted based on the signal-to-noise ratio (SNR), whereas in amplitude-modulation experiments they are varied in modulation depth. When a staircase method is selected (cfg_game.adapt = ‘transformed-up-down’ or ‘weighted-up-down’), additional parameters should be specified, including the step size (cfg_game.start_stepsize), whether the scale is linear or logarithmic (cfg_game.step_resolution), as well as other parameters specific to the type of staircase selected.
* _**user.m**: The user file defines the composition of each trial based on the specified target and background noise, and the experiment configuration. This function is responsible for creating the stimulus that will be presented to the participant. In general, it loads the pre-stored waveforms, adjusts their levels (if required), and combines them according to the experimental variable.

An example of application of this structure to a specific experimental context can be found in Section 6.1. This example illustrates how the four core scripts work together to define and implement an experiment, from initializing the necessary files and configuring the experimental variables, to generating the background noise and stimuli.

When running `fastACI_experiment.m`, the function `fastACI_experiment_init.m` is called and populates the variable ‘cfgcrea’ based on information relevant to the toolbox that is contained in the experimental `*_set.m` and `*_cfg.m` files. Subsequently, the `*_init.m` file is executed generating all the required waveforms. The waveforms are stored in a participant-specific folder (see Section 4). The populated variable cfg_crea (after `*_set.m`, `*_cfg.m`, and `*_init.m`) is stored into the *cfgcrea*_*\*.mat* file (see Fig. 1).

Two optional scripts can also be included in the experiment folder:

* _**dataload.m**: a custom data-loading function specific to the experiment. If provided, this function will be used during data post-processing instead of the default `fastACI_getACI_dataload.m` (see Section 5.1).
* _**instruction.m**: a script displaying customized instructions to participants, replacing the automatically generated message.

Table 2 lists the main parameters available to design an experiment within the fastACI toolbox.

**Table 2.**
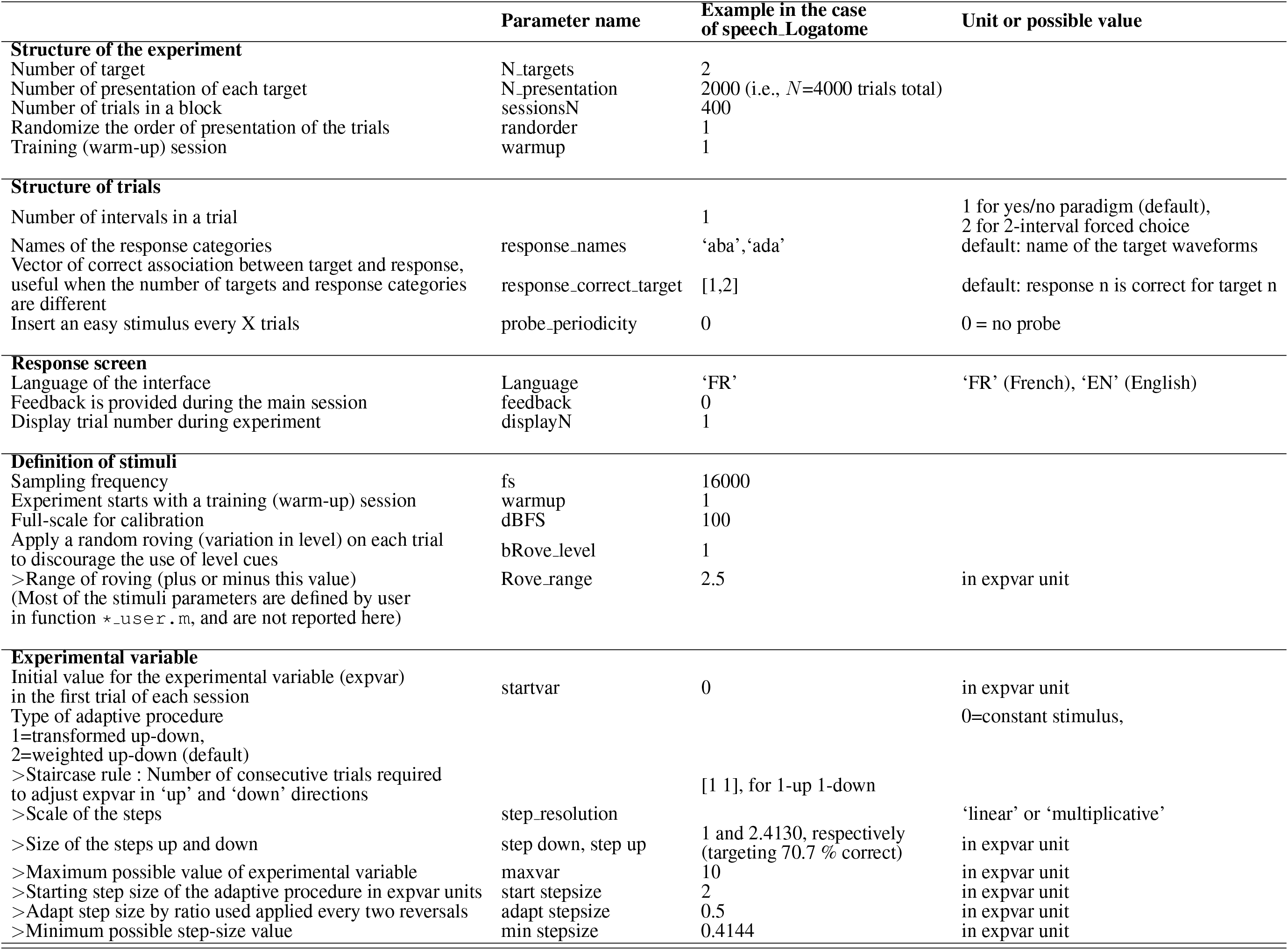
Main parameters for designing experiments in the fastACI toolbox. Symbol ‘*>*’ indicates parameters that depend on the value of a previous parameter.

### 3.4 Running an experiment with an artificial listener

One of our motivations to revisit the old ACI toolbox and convert it into the fastACI toolbox was to enable a listening experiment to be tested not only with human participants but also to replace them by an auditory model. This way, the block “Listener” from the diagram in Figure 1 can actually be toogled to an artificial listener by indicating as subject ID one of the model-reserved words. For instance, using `fastACI_experiment(`‘`speechACI_varnet2013`’,’`king2019`’,’`white`’), i.e., using ‘king 2019’ as a subject ID will automatically run the experiment ‘speechACI_varnet2013’, proceeding with an automatic response simulation of the experiment using the model by King et al. (2019).

Of course the use of reserved words is related to the availability of an auditory model matching that name. We built this artificial-listener mode based on the models available in the AMT toolbox, as of its version 1.0 (Majdak et al., 2022). At the moment of this publication, any of the monaural models available within AMT can be used as an artificial listener by following a couple of steps. We provide now a short and general step-by-step guide in the use of AMT models within the fastACI framework. Given that the post-processing of data collected in an actual listening experiment or using an artificial listener is exactly the same, the next explanation is only focused on getting an auditory model ready for use.

#### 3.4.1 Artificial listener: Front-end auditory model and back-end decision module

Within fastACI, the artificial listeners need to be composed of an auditory front-end module, sometimes referred to as a pre-processing model, and a back-end module that provides a simple binary decision. In such a decision scheme, an incoming sound is labeled as the most likely target interval from a limited set of options, based on signal detection theory. The third-party AMT toolbox provides mainly pre-processing models. Eight of those front-end models have been already comprehensively described in one of our previous studies (Osses et al., 2022b). Despite the fact that the models need to be further configured in order to be successfully used as artificial listeners (see below), if the models exist in the Matlab path, the toolbox will still attempt to use them as artificial listeners. For instance, if ‘king2019’ is indicated, the pre-processing model `king2019.m` will be used.

For the successful use of an auditory model, however, the back-end module providing the binary decision needs to be appropriately configured. Two decision schemes are available in the script `aci_detect.m`. Both decisions schemes are related to the concept of optimal detector (Green and Swets, 1966) with one of the decisions (`cfg_sim.type_decision =` ‘`optimal_detector`’) following the template-matching approach as described by Osses and Kohlrausch (2021) and the other (`cfg_sim.type_decision =` ‘`relanoiborra2019 decision`’) following the decision as used by Relaño-Iborra et al. (2019) in the context of speech tests.

The decision type and other options need to be included in a model configuration file, which has the same name as the auditory model with the suffix ‘*_cfg’ and needs to be visible to Matlab. The expected location within the toolbox is under the folder *Simulations*. A number of configuration files that we have used in previous studies can be found in the folder *Simulations/Stored*_*cfg/*. For the case of the ‘king2019’ model, to use a specific configuration you can copy one of the stored configurations, either `osses2022_02_AABBA_king2019.m` (Osses et al., 2022c) or `osses2023b_FA_king2019.m` (Osses and Varnet, 2023)—both used in speech experiments—and paste it into *Local* as `king2019_cfg.m`. Because these two stored configurations were extensively used by us at the time of the corresponding publications, the artificial listener will be hereafter ready for a successful use within the fastACI toolbox.

#### 3.4.2 Brief explanation of a model configuration script

A model configuration script, called `king2019_cfg.m` in the case of the AMT model `king2019.m` contains sections defining the detectors, defining the template or whatever “expected signal” might be used by a model, extra parameters to be used when calling the model within the third-party AMT toolbox, and finally a compulsory model initialization following the guidelines of the third-party AFC toolbox.

**Listing 1.**
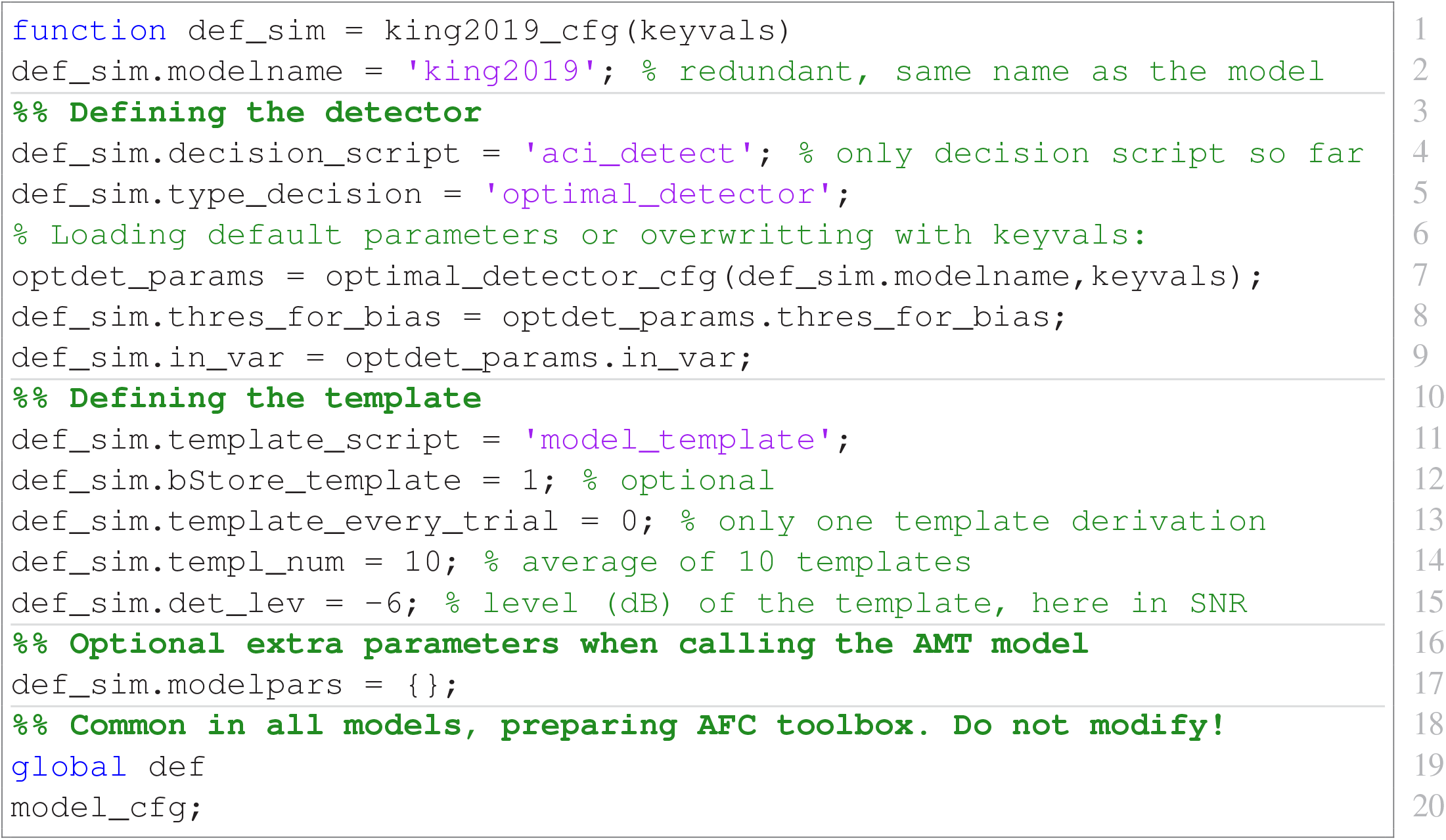
Example of a model configuration file for the front-end model ‘king2019’ using the optimal detector decision (Osses and Kohlrausch, 2021).

The definition of the template should be matched to the relevant parameters of the experiment. From all the list of parameters there, the most critical parameter is the field ‘det lev’ which corresponds to the so-called supra-threshold level, i.e., a value of the dependent variable at which the task will be very easy to solve by the (artificial) listener. In the example of the speech experiment ‘speechACI_varnet2013’, −6 corresponds to a signal-to-noise ratio of −6 dB, at which the ‘king2019’ model was able to solve the speech task nearly perfectly.

Another important field, is ‘modelpars’. In the example above, that field is empty, meaning that ‘king2019’ will only be called using the input signal (the incoming interval sounds, ‘insig’) and the corresponding sampling rate (‘fs’) as input parameters to the model, such that the processed sound ‘outsig’ is obtained from AMT as `outsig = king2019(insig, fs)`. If the user needs to force or change any of the model optional parameters, e.g., specifying `def_sim.modelpars =` {‘`compression_n`’`,0.3`}, then those entries will be appended to the AMT call, resulting in: `outsig = king2019(insig, fs,` ‘`compression_n`’`,0.3)`.

## 4 STORING THE DATA

When the toolbox is run for the first time, the user must specify the location of two compulsory data directories (along with additional folders for dependencies). The first directory, *dir*_*data*, stores all experimental stimuli, while the second one, *dir*_*datapost*, holds all post-processing data. By default, both directories point to the same location, allowing analysis results to be stored alongside raw data for convenience. Setting *dir*_*datapost* to a separate location can be useful if one want to re-generate easily all analysis results from scratch, because in principle all the data to be stored under this directory can be re-generated at any moment using the information contained in *dir*_*data*.

The *dir data* folder follows a hierarchical tree structure: main folder *>* experiment folders *>* participant folders. By default, each participant folder contains two subdirectories: *NoiseStim* which stores waveforms of all noise stimuli presented during the experiment (typically a very large folder) and *Results* which contains the *cfgcrea*_*\*.mat* and *savegame*_*\*.mat* files. If the experiment involves complex targets that cannot be entirely defined within the user function, such as in speech perception tasks, a *speech-samples* subfolder is also included in the participant directory.

If *dir*_*datapost* is set to the same location as *dir*_*data*, an additional *Results_ACI* subfolder appears within each participant’s *Results* folder, storing post-processed data derived from the savegame files, including the computed ACI.

The *cfgcrea*_*\*.mat* and *savegame*_*\*.mat* files, stored in the *Results* folder, are generated at different stages in the experiment and contain partly redundant information about the experiment initialization, and the data collection. The final file name contains information such as the participant ID, the time stamp of file creation, and the experimental condition, if relevant. We provide now more details about this information.

### 4.1 *cfgcrea_*.mat* file: cfg_crea and info_toolbox structures

The initialization file, *cfgcrea*_*\*.mat*, contains two struct variables cfg_crea and info_toolbox, the latter storing information about the toolbox version. The variable cfg_crea contains compulsory and optional fields describing the experiment. Some of these fields correspond to information provided in the experiment files (see Section 3.3), while others are automatically obtained during initialization, such as stim order which defines the actual presentation order of the trials. The information contained in cfg_crea is then passed to the cfg_game structure.

### 4.2 *savegame *.mat* file: cfg_game and data_passation structures

Each time a session concludes, the experiment ends, or the participant requests a break, a new *savegame*_*\*.mat* file is created and stored in the participant’s *Results* folder. To avoid accidental data loss, previous savegame files are not deleted automatically but moved to a different subfolder *Results*_*past*_*sessions*. However, since each newly created savegame file contains all information recorded in the earlier ones, it is possible to manually remove previous files without losing any data.

The *savegame*_*\*.mat* file contains two struct variables, cfg_game and data_passation. The cfg_game variable contains all the fields from cfg_crea, in a way that no later access to cfg_crea is needed to post-process the collected experimental data. Additionally, cfg_game contains information about the collected data, in particular the responses of the participant. As for all trial-specific variables in cfg_game, the responses are stored in the order of the noise in the *NoiseStim* folder, which does not necessarily correspond to presentation order.

The *savegame*_*\*.mat* file also contains a variable called data_passation, that contains all the data that are being (or was already) collected. This is the structure that will be processed in later analysis stages. For this reason, the trial-by-trial data in data_passation is arranged based on the order of presentation (stored in cfg_game.stim_order), rather than in alphabetical order as in cfg_game and cfg_crea, making it easier to analyze and plot data temporally. A schematic overview of the data organization for a five-trial experiment is shown in Figure 2.

**Figure 2.**
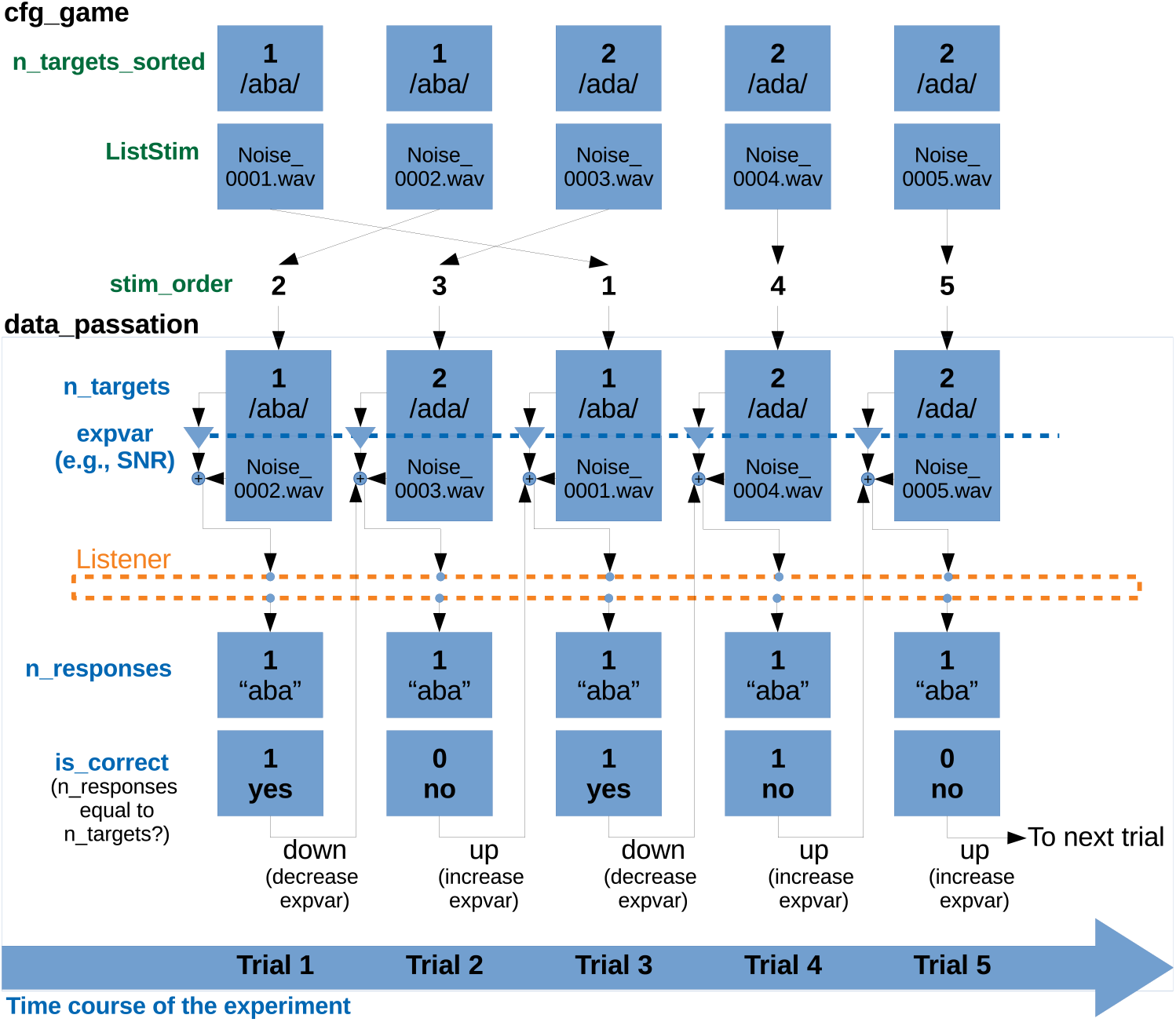
Schematic representation of the relevant fields in the variables cfg_game and data_passation, for a 5-trial experiment following a simple 1-interval 2-alternative paradigm with a 1-up 1-down staircase procedure. The variables in cfg_game (here, n_targets sorted and ListStim) are sorted by the number of the corresponding noise file. The variable cfg_game.stim_order defines the actual presentation order. All variables contained in data_passation are sorted according to presentation order. Therefore, cfg_game.n_targets_sorted and data_passation.n_targets represent the same information, sorted in different ways. data_passation also contains the tracking variable that is stored as expvar (here, the SNR) and the participants’ responses. The variable data_passation.is correct is obtained by comparing of n_targets and n_responses. If the response is correct, the tracking variable will be set to a down run (to a more difficult condition) and if the response is incorrect, to an up run (to an easier condition). [Updated: wavefiles numbered from 0001–0005; expvar is SNR; replace /ba/ with /aba/; add “time course of the experiment”]

### 4.3 *ACI_*.mat* file: cfg ACI and results structure

Finally, the post-processing of the data, described in Section 5 results in a third type of .mat file, the *ACI_*.mat* files. Unless specified otherwise, they are stored in a *Results_ACI* folder within the corresponding participant-specific *Results* folder and are labeled using the following naming convention: `ACI-SID-expName-cond-trialtype-dataload-fitting_function-last-expvar`. In this naming scheme, `SID` refers to the participant’s identifier, `expName` to the experiment name, and `cond` to the experimental condition, which might be empty if the experiment only has one condition. The next labels correspond to the first three stages of the data post-processing: `trialtype` refers to the type of trials used for analysis (Section 5.2), `dataload` is a short identifier for the data-loading function (Section 5.1), `fitting_function` is a short name for the fitting function to be used (Section 5.3), `last` indicates the number of the last trial—relevant if the experiment has not been fully completed yet—, and `expvar` a short name for the trial selection criterion based on dependent “expvar” variable (Section 5.2). Although this naming convention covers the main choices experimenters have to make when calculating an ACI, it is not precise enough to distinguish all possible processing pipelines. This is why the `fastACI_getACI.m` function, which computes and stores the ACI, also includes an option *dir*_*out* that allows the experimenter to indicate a different folder for storing the resulting mat file and/or add a prefix to the name.

As for the previous *cfgcrea*_*\*.mat* and *savegame*_*\*.mat* files, the *ACI*_*\*.mat* file contains two data structures. cfg ACI lists all options selected for the computation of the ACI, as well as the version of the toolbox used. The results structure contains all outcome measures, in particular the estimated ACI and its dimentions. Depending on the options used for the computation (see Section 5.3), it can also include information about the fitting process such as the hyperparameter values tested and about the validation of the final ACI (see Section 5.4). For convenience, the final outcome of the estimation process is also stored as a matrix variable named ACI.

### 4.4 Recreating the noise waveforms

Revcorr experiments typically require a substancial memory space, as a unique set of noise is generated for each participant, and every waveform must be stored on disk during data analysis. For example, in the study of Carranante et al. (2024), 49 datasets were collected, each of which contains 4000 noise stimuli. With each waveform being 27.3 kB, the total storage required amounted to 5.35 GB. One important feature of the toolbox is the possibility to store only the random seeds used to generate the noisy stimuli, rather than the stimuli themselves. The seeds are saved in the *cfgcrea*_*\*.mat* file during the initialization phase. After the data collection is completed, the experimenter can delete all stimuli waveforms from disk, and re-generate them when needed. In Carranante et al. (2024)’s study, each *cfgcrea*_*\*.mat* file is 29.3 kB, reducing the total storage requirement to less than 1.5 MB.

For any experiment, the sound waveforms used by a participant can be retrieved from the stored cfg_crea files using the command: `fastACI_experiment_init_from_cfg_crea(`‘`cfg_crea_name`’). This function first checks if the corresponding *NoiseStim* directory within the participant’s folder is empty. In such a case the function calls the experiment-specific `*_init.m` function which re-generates the noise set. Alternatively, the user can directly call the `*_init.m` function with the cfg_crea or cfg_game variable as argument.

## 5 POST-PROCESSING OF THE DATA

The fastACI toolbox offers the possibility to post-process the collected data through a revecorr analysis. This aims at finding a statistical relationship between the random stimulus presented in each trial and the corresponding response of the participant. As for experiment design (Section 3) our objective was to make this module as versatile as possible, in particular allowing different types of signal representation and the fitting of different statistical models.

The data post-processing results in a participant-specific matrix of weights that reflects the influence of the random fluctuations in the signal on the participant’s response. These matrices are often referred to in the visual perception literature as “classification images.” For this reason, we usually refer to the output of the auditory revcorr experiments as auditory classification images (ACIs) (Varnet et al., 2013). Other names in the literature notably include “participant weightings” (Ahumada et al., 1975), or “kernels” (Varnet and Lorenzi, 2022; Joosten et al., 2016).

The central function of the post-processing module is the script `fastACI_getACI.m`. This script performs the following processes in sequential order: (1) it loads all waveforms (secondary script `fastACI_getACI_dataload.m`), (2) it selects the specific trials to be further processed and applies transformations to the data if needed (secondary script `fastACI_getACI_preprocess.m`), (3) it computes the ACIs (secondary script `fastACI_getACI_calculate.m`).

The `fastACI_getACI.m` script requires as input the binary savegame file, from where the variables cfg_game and data_passation are loaded. If no other arguments are included, default parameter values are used, corresponding to the simplest analysis pipeline: a correlation analysis based on gammatone spectrogram representations. Optional parameters can be transmitted as additional input arguments. The main options for computing an ACI are listed in Table 3.

**Table 3.**
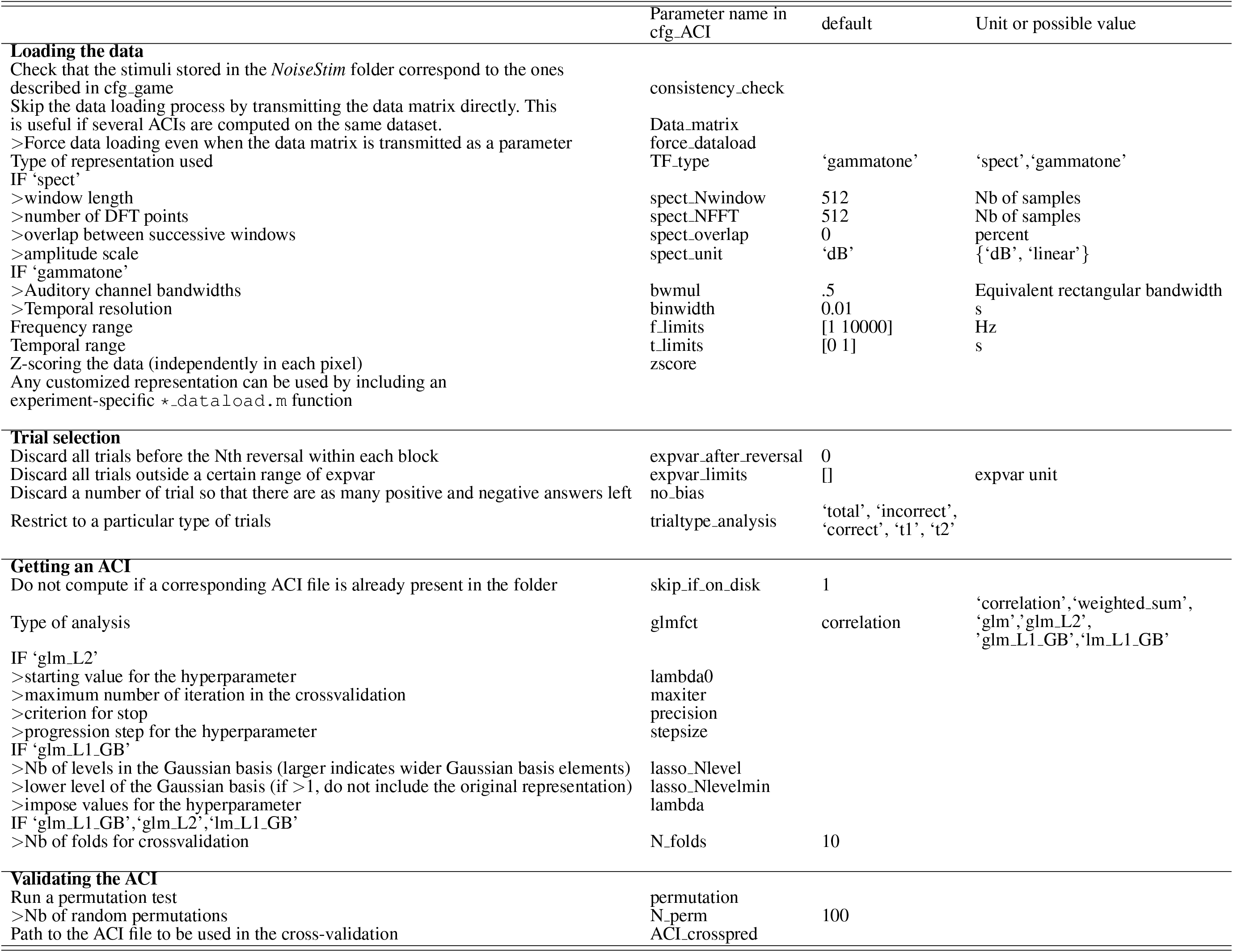
Main parameters for post-processing the data in the fastACI toolbox. Symbol ‘*>*’ indicates parameters that depend on the value of a previous parameter.

### 5.1 Stage 1. Loading the data: fastACI_getACI_dataload.m

The function that loads the data plays a critical role, as it reads the stimulus waveforms and converts them into a matrix that is subsequently used for the ACI assessment. The dimensions of this matrix determine those of the resulting ACI, which are identical. The default script for data loading is `fastACI_dataload.m` but it can be overridden if an experiment-specific function named `⋆_dataload.m` is found on disk (see Section 3). An example of such a function can be found for experiment `modulationACI_dataload.m`.

The dataload function returns a matrix containing the (typically time-frequency) representations of all noise instances. By default, the first dimension corresponds to the trial number, in ascending order of presentation, while the second and third dimensions are related to time and frequency, respectively. The default representation is a time-frequency gammatone-based spectrogram, obtained through the `Gammatone_proc.m` function. This representation has a temporal resolution of 0.01 s and a spectral respolution of 0.5 in the Equivalent Rectangular Bandwidth Number scale (ERB_*N*_). The frequency dimension is obtained from a critical filter bank covering centre frequencies between 45.8 Hz (1.69 ERB_*N*_) and 8000 Hz (33.19 ERB_*N*_). This results in 64 frequency “bins,” followed by an envelope extractor based on a simplified inner-hair-cell processing (Osses et al., 2022b).

Although time-frequency representations, like the one described above, offer an intuitive way of interpreting the perceptual weights, the second and third dimensions can contain any alternative stimulus feature estimate. The third dimension is optional; if it is not specified, the obtained ACIs will only be two-dimensional. Using an experiment-specific data-load function can be useful if the experimenter is interested in exploring specific dimensions of the stimuli. There are two situations where this option might be particularly relevant:

- First, if the experiment involves background noise covering the entire time-frequency space, but the experimenters are only interested in a specific acoustic feature, it can be beneficial to perform the revcorr analysis on this dimension alone. This is the approach followed in experiment `modulationACI` (Varnet and Lorenzi, 2022), where the experiment focused on the role of the envelope in a single frequency band. In this case, the experimenters defined an experiment-specific data-load function (`modulationACI_dataload.m`) to represent only the envelope in the selected frequency band.
- Second, if the experiment is based on a customized `*_user.m` function instead of the default `Generate_noise.m`, defining the data-load function accordingly is advisable. A good example of this is the prosodic revcorr experiment, where the targets are not embedded in a background noise but are resynthesized with a random prosody (Osses et al., 2023, experiment `segmentation`). In this case, the `segmentation_user.m` function generates the stimuli and stores the trial-by-trial parameters of the random prosody, while the `segmentation_dataload.m` function loads these parameters and organizes them into a 2-dimensional data matrix.

### 5.2 Stage 2. Selecting the trials: fastACI_getACI_preprocess.m

This pre-processing step optionally prepares the matrix obtained from the data-loading function before it is analyzed by the `fastACI_getACI_calculate.m` function. The primary role of this stage is trial selection. Although the default option is to bypass this step and conduct the revcorr analysis on all trials, there are situations where it is beneficial to exclude specific trials prior to further analysis. Trial selection is controlled by four parameters that can be fully combined:

#### ‘trialtype_analysis’

This parameters allows for the selection of specific trials. By default, the ACI is computed across all trials using the parameter value ‘total.’ However, separate calculations for target-present and target-absent trials can provide valuable insights into the influence of nonlinear auditory processing (Ahumada et al., 1975). In particular, the target-absent ACI is considered a better estimate of the true underlying internal or “mental” template of the participant in the presence of non-linearities in the processing. Such target-specific analyses can be carried with parameter values ‘t1’ and ‘t2’, selecting the trials depending on the number of the target that was presented (1 or 2, respectively). Alternatively, it may be useful to restrict the analysis to correct or incorrect trials only, as was done by Osses and Varnet (2024). This can be achieved by specifying the parameter values ‘incorrect’ or ‘correct’, respectively.

#### ‘expvar after reversal’

This parameter controls the exclusion of initial trials in a staircase procedure. In an adaptive experiment, the trials at the beginning of each block correspond to the convergence of the staircase, transitioning from the initial expvar value (startvar) to the perceptual threshold defined by the staircase rules (see Section 3.3). However, these early trials typically provide little information for the ACI but tend to introduce noise into the estimation, as expvar can take very large or very low values. For this reason, they are often rejected from analysis. Any value larger than zero for the ‘expvar after reversal’ parameter specifies the number of staircase reversals to exclude from further analysis.

#### ‘expvar limits’

Similarly, it can be useful to discard trials corresponding to extreme expvar values throughout the experiment. For instance, if expvar correspond to the SNR in dB at which targets are presented, very low expvar values correspond to trials where the target is virtually inaudible and the participant may respond at random. Therefore, these trials do not provide any valuable information on the underlying auditory mechanisms. Conversely, very high expvar values correspond to easy trials where the noise has no impact on the decision. As such, these trials do not contribute either to the ACI estimation and removing them typically results in more reliable estimates.

#### ‘bias’ / ‘no_bias’

This last parameter explicitly controls for the balance between the two types of responses. If the flag ‘no bias’ is included, a number of trials will be selectively removed to balance the responses. In other words, if a participant indicated “response 1” 53% of the times and “response 2” 47% of the times, a number of “response 1” trials, corresponding to 6% of the total, will be excluded. For reasons mentioned above, trials are discarded based on the absolute distance from the mean expvar, so that the trials with the most extreme expvar values are excluded first.

Another optional pre-processing step is controlled by the parameter ‘zscore.’ When this parameter is set to 1 (the default), the data matrix after trial exclusion is z-scored independently for each element in the representation (that is, along the first dimension only). In the context of the typical target-in-noise task, this option is especially useful when the noise energy does not have the same variability across every time-frequency pixel (see Figure 3). Z-scoring normalizes the ACI weights, allowing them to be interpreted as perceptual weights. If the experimenter has specified a custom `*_user.m` function, z-scoring can also be useful as it allows expressing all ACI weights on a consistent, unitless scale. This normalization facilitates comparison across different dimensions of the stimulus.

**Figure 3.**
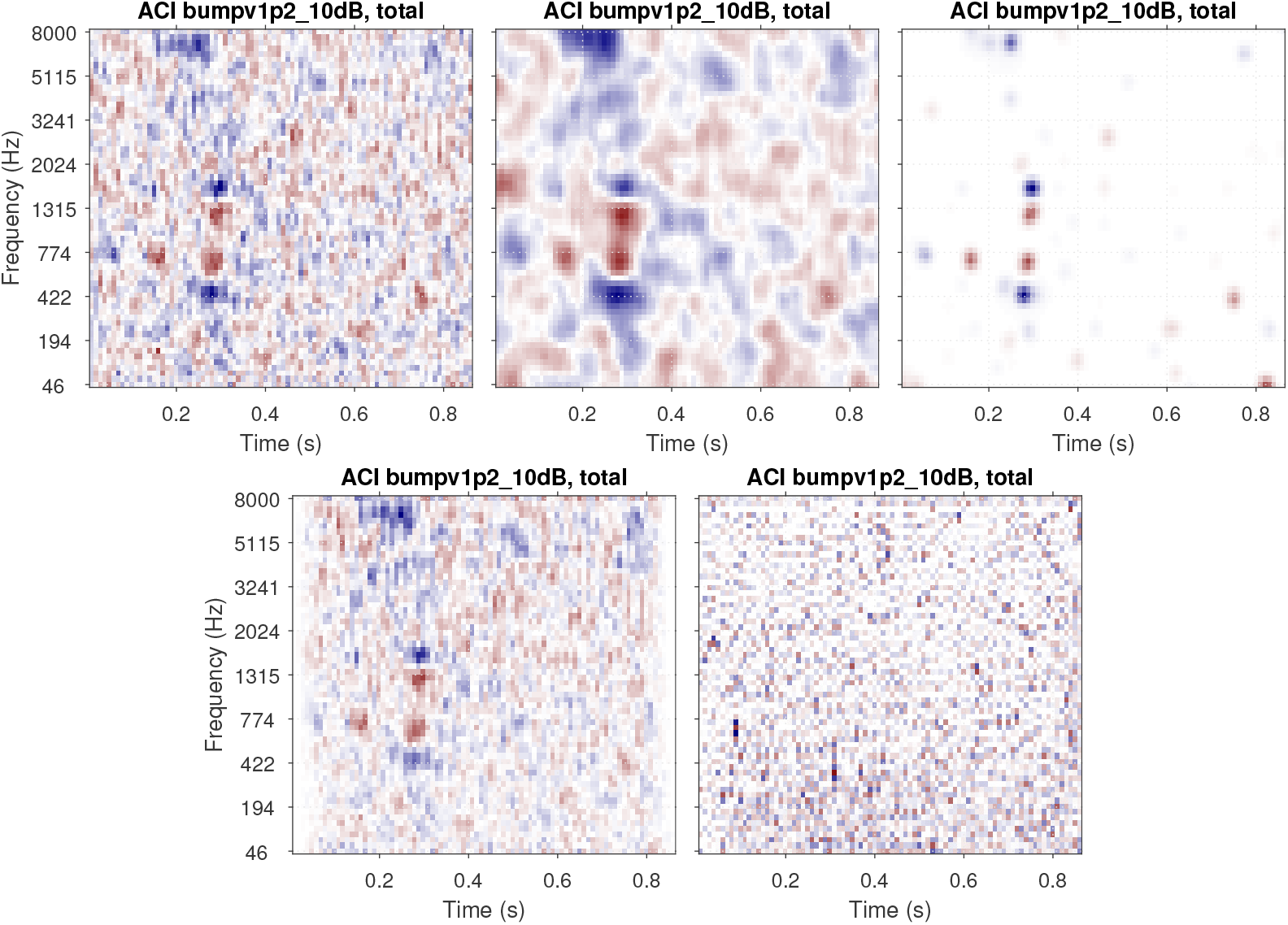
ACIs derived from a single dataset (participant S04 from Osses and Varnet (2024), 4000 trials of aba-ada categorization in bump noise) analyzed using different algorithms. The top row shows the recommended approaches: correlation (left), GLM with L2 norm penality (center), GLM with L1 norm penality on a Gaussian Basis (right). The bottom row presents approaches that do not yield easily interpretable ACIs in general: weighted sum without z-scoring (left) and GLM with Maximum Likelihood estimation (right). Apart from the type of analysis, all parameters are set to their default value. All ACIs are normalized in maximum absolute weight. [color bars removed]

### 5.3 Stage 3. Getting an ACI: fastACI_getACI_calculate.m

In this critical stage, the pre-processed data matrix from the previous step is analyzed together with the response vector, to examine the influence of noise on perception. This is achieved through a revecorr approach, which identifies the statistical relationship between the random fluctuations of the noise presented in a given trial and the corresponding binary response of the listener (“target 1” or “target 2”). The outcome of this analysis is summarized as an auditory classification image (ACI) with the same dimensions as the stimulus representation chosen in the previous stage (typically, time-frequency).

More specifically, the ACI analysis allows to identify which features in the noise bias the decision of the listener towards one alternative or another. In other words, this computation highlights the (typically time-frequency) regions of the stimulus that the listener relies on as cues for resolving the task. The ACI represents these cues by associating individual weights to each pixel in the noise representation, quantifying how much each particular element contributes to the final decision. These weights are often interpreted as estimates of the “perceptual weights” the participant attaches to each acoustic features, while the ACI is sometimes considered as a visualization of the internal or “mental” representation of the target sounds, that are formed, stored and used by the participants. A discussion of the limitations of these interpretations is beyond the scope of this paper. Suffice it to say that the analysis itself does not rely on any assumption about the existence or nature of any perceptual weights or internal representation. As Neri (2018) argued, classification images can be regarded as a descriptive statistics summarizing the data, much like the mean or the median, rather than as estimates of underlying perceptual components.

The function `fastACI_getACI_calculate.m` handles the revcorr analysis. Currently, the toolbox offers five main computational options for this stage, each based on a different statistical model: ‘correlation’, ‘weighted sum’, ‘glm’, ‘glm_L1_GB’, and ‘glm_L2’. Each of them takes as input the noise matrix 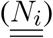 and the behavioral responses (*r*_*i*_) and return an ACI matrix 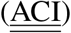. With *N* _trial_ the number of selected trials for the analysis, *N*_*f*_ the number of bins for the first dimension of the stimulus representation (typically, frequency), and *N*_*t*_ the number of bins for the second dimension of the stimulus representation (typically, time), 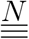 is a *N*_*trial*_-by-*N*_*f*_ -by-*N*_*t*_ matrix, *<u>r</u>* is a binary vector of length *N* _trial_, and 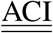 is a *N*_*f*_ -by-*N*_*t*_ matrix. In the following, we will denote <u>ACI</u> the ACI matrix in its vector form (i.e. a *N*_*f*_ *× N*_*t*_-by-1 vector) and *<u>N</u>*_*<u>i</u>*_ the vectorization of the noise matrix for trial *i*.

The next sections present the mathematical framework for the five main options for computing an ACI, as well as their limitations. The result of these different estimation methods applied on a single set of data are shown in Figure 3. In Section 6.5, the different options are compared with regards to the goodness of the fit.

#### 5.3.1 Correlation and weighted sum

A straightforward and intuitive way to summarize the relationship between stimuli and participant’s responses is to compute their correlation. In this case, the value of the ACI in each time-frequency pixel *j*, denoted as ACI_*j*_, is simply given by the Pearson correlation coefficient between the corresponding pixel in the stimuli representation *N*_*i,j*_ and the vector of responses *r*_*i*_ across all trials *i*. This method is implemented through the ‘correlation’ option.

Another very common approach is the so-called “weighted sum” ACI, which is calculated by subtracting the average noise representation for “response 2” from the average noise representation for “response 1.” This method is available via the ‘weighted sum’ option. It can be shown that the two options are equivalent up to a multiplicative factor, under the assumptions that the noise is centered with a constant variance (this is ensured if the ‘zscore’ option is enabled) and that the participant is unbiased, i.e., *P* (*r*_*i*_ = “response 1”) = *P* (*r*_*i*_ = “response 2”) = 0.5.

Because of their simplicity, these two options are fast to compute. Furthermore, they only rely on the general assumptions, common to all revcorr approaches, that cue detection is influenced by random fluctuations introduced in the stimuli and that cues are confined to the dimensions of the representation. Importantly, these methods do not make assumptions about the specific shape of the cues. However, a downside of these approaches is that the resulting ACI is often relatively noisy due to overfitting: when the number of predictors is relatively large compared to the number of trials, the ACI can capture spurious correlations in the noisy data, which may obscure the relevant features.

An example of ACI obtained through the correlation procedure is shown in Figure 3 (top left), which results in an accurate (although noisy) ACI, with larger positive and negative weights in the regions corresponding to the cues. The weighted sum approach with z-scoring yields an identical result. The bottom left panel of Figure 3 shows an ACI computed using the weighted sum approach without z-scoring. Due to the non-uniform distribution of noise across the time-frequency space, caused, for example, by the use of fade-in and fade-out ramps in this experiment, this second ACI displays a distinct weight pattern. Here, the weights reflect both the participant’s responses and aspects of the noise’s statistical distribution. In particular, the smaller variability at stimulus onset and offset, yields smaller weights in these regions, regardless of whether the information is actually used by the participant. This approach can complicate the interpretation of the ACI, as it becomes difficult to disentangle whether a given weight reflects the statistics of the stimulus or the participant’s response, and it should therefore generally be avoided.

#### 5.3.2 Linear regression

Assuming that the variance of the noise is the same in each pixel (true if the ‘zscore’ option is enabled) the previous ACI approaches are equivalent, up to a multiplicative factor, to performing independent linear regressions on each pixel *j*:

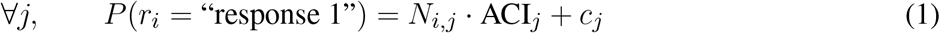

with ACI_*j*_ and *c*_*j*_ corresponding to the regression coefficient, fitted by maximum likelihood.

This naturally suggests gathering all predictors within a single linear model, as in Ahumada et al. (1975):

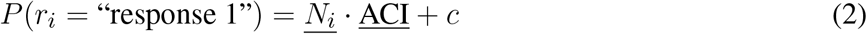

This multiple linear regression approach is rarely used as it generally suffers from three issues: overfitting (as highlighted above), multicollinearity, and heteroscedasticity.

Multicollinearity and overfitting will be discussed in the next sections. Heteroscedasticity refers to non-uniformly distribution of prediction errors. In the case of the models above, this is evident from the fact that, while the left-hand member is a probability bounded between 0 and 1, the right-hand member can theoretically vary from -∞ to +∞. Although this does not necessarily pose a problem in practice, as the probabilities rarely approach floor and ceiling values in a revcorr experiment, this incompatibility of the distributions described by the two members of the equation prompts us to look for a more adequate model.

#### 5.3.3 Generalized linear model with maximum likelihood estimation

A natural solution, introduced by Knoblauch and Maloney (2008) and implemented within the toolbox (‘glm’ option), consists in replacing the linear regression by a generalized linear model (GLM). As the dependent variable (the response of the participant) follows a binomial distribution, it is better modeled through a normal cumulative distribution function Φ, linking the linear combination of predictors to the probability *P* (*r*_*i*_ = “response 1”):

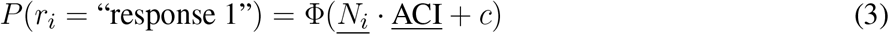

The approach described in Equation (3) is particularly useful when the experiment has a limited number of predictors relative to the number of trials, and each of these predictors are statistically independent one of each other. This is for instance the case in the study by Osses et al. (2023), for which their statistical model had 16 variables (each corresponding to an independent Gaussian distribution), that was individually fitted to 800 observations of each participant. In the general case, however, these conditions may not be met, and the model will lead to highly noisy ACIs. This is the case for instance when this approach is applied to the data of Osses and Varnet (2024) (Figure 3, bottom right panel), as there is a large number of predictors (*N* _trials_ = 4000 and each noise is described using 5504 time-frequency bins), which are highly correlated to each other. These two factors can give rise to overfitting and multicollinearity issues, respectively, each of which compromising the accuracy of the estimation.

Overfitting arises when the number of predictors is large relative to the number of observations and results in a lack of generalization ability. This is because, in this case, the statistical model—this GLM—is able to capture not only the meaningful patterns in the data but also spurious correlation. Multicollinearity refers to the presence of correlations between predictors within a statistical model. It can lead to counter-intuitive results where none of the predictors appear to be directly related to the dependent variable, even though, in reality, all predictors are associated with it.^2^ Multicollinearity can therefore lead to a severe underestimation of the ACI. However, if the predictors in Equation (2) are chosen to be statistically independent of each other, multicollinearity is no longer an issue, and this approach becomes equivalent to the previous one, from Equation (1). Note that the independent linear regression approach of Equation (1), as well as the equivalent correlation and weighted sum approaches, are immune to multicollinearity as each predictor enters a separate statistical model.

#### 5.3.4 Generalized linear model with regularizers

The fitting of statistical models presented in the previous sections, Equations (2) and (3), is most often performed using maximum likelihood estimation. However, as we have pointed out, this approach can lead to inconsistent parameter values or imprecise estimates. One possible solution to address both overfitting and multicollinearity in regression is by introducing a regularizing prior, through penalized regression. In the context of classification images, this solution was proposed by Knoblauch and Maloney (2008) and later adopted by Mineault et al. (2009). Two regularizing priors are currently implemented in the toolbox: L2 regularization (‘glmfitqp’ option) and L1 regularization on a Gaussian basis (‘glm_L1_GB’ option). A detailed description of each of these priors can be found in the studies by Varnet et al. (2013) and Osses and Varnet (2024), respectively. Here, we summarize the general framework of penalized regression.

As the name indicates, the maximum likelihood approach identifies the parameter values (<u>ACI</u> and *c*) that maximize the likelihood *L*({<u>ACI</u>; *c*}) given the observations. This is equivalent to minimizing the negative logarithm of the likelihood, also known as the negative log-likelihood. Penalized regression consists in minimizing both the negative log-likelihood and an additional penalty term *P* ({<u>ACI</u>; *c*}), which also depends on the parameters. The relative weight assigned to these two terms is determined by a hyperparameter *λ*, leading to the following minimization objective:

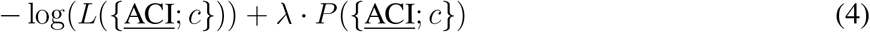

The penalty term reflects prior knowledge about the plausible values of the parameters. For example, Varnet et al. (2013) and Varnet et al. (2015a) employed a smoothing penalty (L2 regularization) based on the assumption that the estimated ACI should not exhibit abrupt discontinuities—a relatively natural assumption given the spectral and temporal resolution of the human auditory system. Conversely, the approach followed by Osses and Varnet (2024) and Carranante et al. (2024) is based on a lasso penalty (L1 regularization) applied to a Gaussian basis. This approach assumes that most ACI weights are zero, except in specific regions with a Gaussian shape in the time-frequency space. As these regularizers implement slightly different assumptions, they result in different ACIs (see Figure 3, top row, center and right panels). The selection of a specific prior is therefore critical and should be informed by our understanding of the perceptual processes involved (Mineault et al., 2009). In the case of consonant perception, for instance, it is well-established that listeners rely on acoustic cues that are highly localized in both time and frequency. Therefore, L1 regularization on a Gaussian basis is advisable in this case, and yields better estimates (see Section 6.5).

The relative importance of the regularization and the likelihood (i.e., between the data and the a priori knowledge injected into the estimation) is controlled by the hyperparameter *λ* in Equation (4). As the name hyperparameter indicates, *λ* is not a parameter of the statistical model of Equation (3), but of the estimation itself. Any predetermined hyperparameter value will result in a particular fit of the model with a particular influence of the regularizer: high *λ* estimates are exaggeratedly distorted by the regularization, while the solution approaches that of maximum likelihood when *λ* approaches zero. As represented in Figure 4, an intermediate *λ* value (here *λ* = 0.024) corresponds to a realistic estimate. The greater reliability of the corresponding ACI can be quantified by its out-of-sample predictive accuracy: typically, the ability to predict new data is low for small values of lambda, as overfitting would lead to poor model generalizability. For very large *λ* values, the penalty term becomes predominant over the data, resulting in a decline in predictive quality. Out-of-sample predictive accuracy is assessed in terms of cross-validated deviance (see Section 5.4.3).

**Figure 4.**
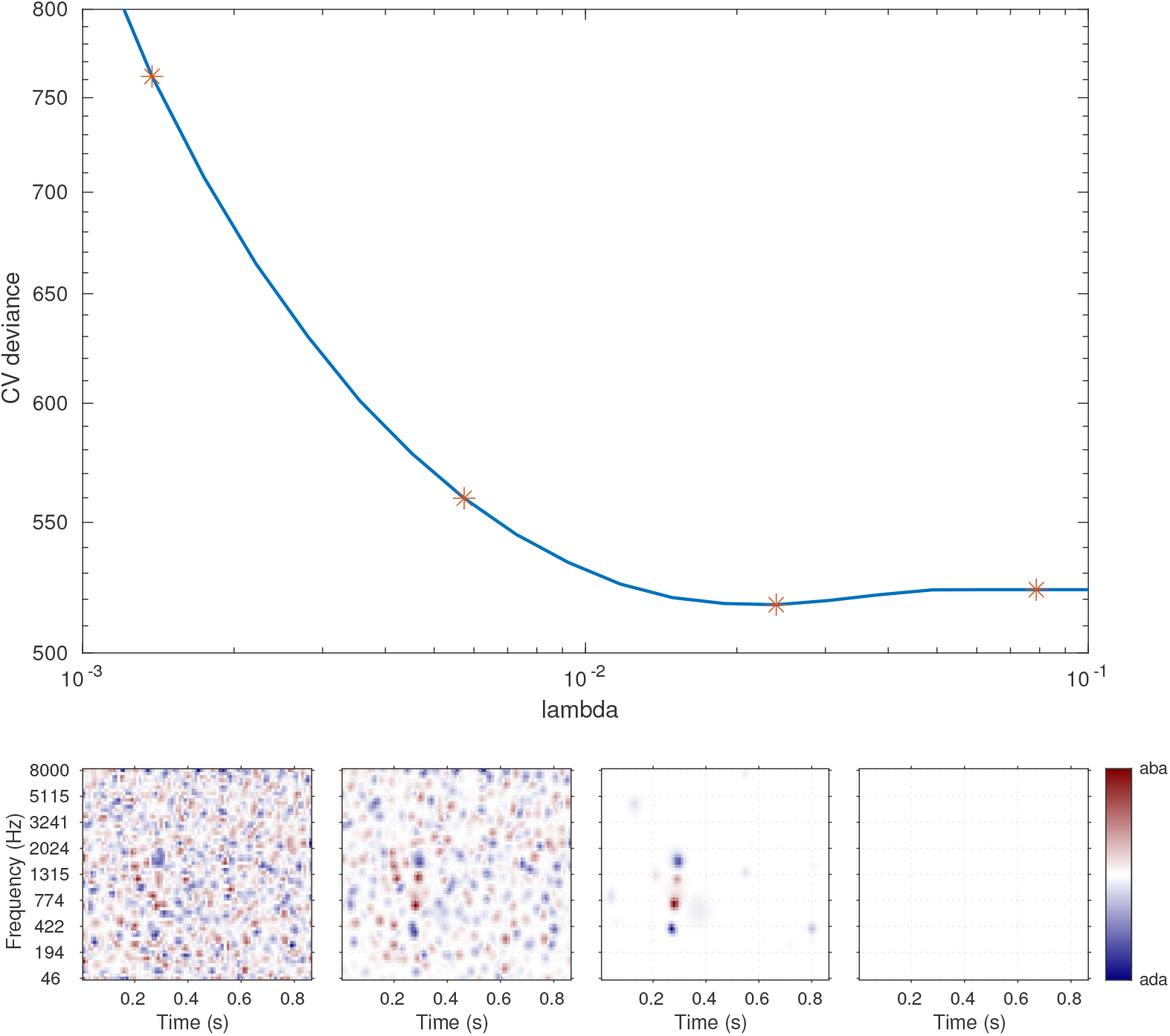
Illustration of the hyperparameter selection process, based on the data from participant S01 in the MPSN condition in Osses and Varnet (2024). Top panel: Cross-validated deviance as a function of the value of the hyperparameter *λ* used for the fit. Bottom panels: ACIs estimated using four hyperparameter *λ* values (indicated by stars in the top panel). The third ACI corresponds to the optimal hyperparameter value (here, *λ* = 0.024).

Penalized regression adresses both the multicollinearity and overfitting issue. It recognizes the presence of dependencies in the prediction and explicitly uses them in the fitting process to reduce the number of effective predictors, thus lowering the risk of multicollinearity. Moreover, as explained above, the hyperparameter selection criterion is based on the ability of the GLM to generalize, protecting the estimation against potential overfitting effects.

Regardless of the estimation option selected, the `fastACI_getACI_calculate.m` function handles the fitting process, returning the final ACI, along with any relevant variables computed during the estimation. This script chooses the final ACI as the one that has the *λ* value that minimizes the cross-validated deviance. Several optional parameters can be specified, including, e.g., the range of *λ* values considered (see Table 3).

### 5.4 Stage 4. Validating the ACI

Regardless of the statistical framework used for the estimation, ACIs inherently involve some amount of estimation error, complicating the interpretation of the results. In the context of an experimental study, it is crucial to conduct statistical validation on the obtained images, to assess whether the ACI genuinely reflects the listener’s strategy or is simply the result of estimation noise. In the following paragraphs we describe several statistical methods implemented within the toolbox. Although they are referred to as a separate post-processing stages, the computations are often nested with those of Stage 3.

There are two primary types of validation methods available: (1) global validation of the ACI, by evaluating whether the underlying model can reliably predict new data from the same participant or from another, and (2) validation of specific weights in the ACI to determine if a cue is present at a particular time-frequency location, using regression coefficient statistics or a model-independent permutation test. No correction for multiple testing is applied to the weight-specific statistics, because such corrections are inherently linked to the specific hypotheses being tested. However, unless the experimenters are interested only in the significance a single predefined weight or group of weights, it is recommended that they implement an appropriate form of multiple testing correction adapted to their needs.

The results of the validation process are returned alongside the estimated ACI itself, as described in Section 4.

#### 5.4.1 Regression coefficient statistics

Most estimation approaches described in Section 5.3 rely on a specific statistical model (linear model for correlation approach, generalized linear model for all GLM-based approaches). When fitting these models, the procedure does not only estimate the optimal weights for the ACI but also calculates the corresponding test statistics and p-values, which reflect the significance of each weight. These statistics are automatically included in the ACI output, providing information about the reliability of the estimated weights.

#### 5.4.2 Permutation test

A common way of assessing which weights in the ACI are large enough to be considered significantly different from zero is through a permutation test. This procedure provides a way to compare the observed weights to a distribution of weights generated under the null hypothesis (i.e., assuming random responses from the participant). The procedure involves generating a large number of random permutations of the participant’s responses (typically 100 permutations or more), and computing an ACI for each of these permutations. The resulting distribution of weights under the null hypothesis is then compared to the observed ACI, allowing experimenters to determine which weights are significantly different from zero.

Although the permutation test can theoretically be combined with any of the ACI estimation methods described in Section 5.3, it can become computationally expensive, especially when using GLM-based approaches.

In the toolbox, the computation of the permutation test can be requested through the flag ‘permutation,’ together with an optional parameter ‘N perm’ indicating the number of permutations (default: 100). The procedure generates *N* _perm_ new datasets by randomly permuting the order of the responses in the original dataset, then computes an ACI for each of these permuted datasets. For each pixel, the 5-th and 95-th percentiles are calculated. The final output includes the *N* _perm_ new ACIs as well as the 90% confidence interval, providing a robust measure of which weights can be confidently attributed to the participant’s response.

#### 5.4.3 Within-participant cross-validation

The two statistical validation methods described above are meant to identify significant regions in the ACI. However, it can also be useful to assess whether the ACI as a whole can be considered a good representation of the participant’s listening strategy in the task. This is usually performed by measuring the ability of the ACI to predict new data from the same participant, using cross-validation. Note that this validation step and the following are for the moment limited to the ‘glm_L1_GB’ option, but they should be extended to ‘glm_L2_GB’,’correlation’ and ‘weighted sum’ in a following release.

Two measures of goodness of fit are computed in the toolbox: the prediction accuracy and the deviance (Osses and Varnet, 2024). Prediction accuracy corresponds to the percentage of correctly predicted binary answers, considering that the model described in Equation (3) responds “1” if *P* (*r*_*i*_ = “response 1”) *≥* 0.5 and responds “2” otherwise. Prediction accuracy is an intuitive metric, but it is usually less precise than deviance. Deviance is the standard goodness-of-fit measure for GLMs, directly related to the log-likelihood.

When the same set of data is used to train and evaluate the model, the goodness of fit is usually overestimated, due to overfitting (see Section 5.3). A solution to obtain an unbiased measure of prediction performance is cross-validation. During cross-validation, the dataset is divided into *N* _fold_ disjoint subsets of equal size. *N* _fold_ − 1 of these subsets are used to derive an ACI, whose prediction accuracy and deviance is evaluated on the remaining subset. The same procedure is repeated *N* _fold_ times to ensure each subsets is used once as the validation set. In this way, the model is never tested on the same trial used for training. The measures obtained for these *N* _fold_ model fits can then be averaged to obtain the average cross-validated prediction accuracy and cross-validated deviance. Furthermore, the dispersion of the *N* _fold_ cross-validated metrics can be used to summarize the reliability of the estimation procedure as a confidence interval (Osses and Varnet, 2024).

#### 5.4.4 Between-participant cross-validation

The above goodness-of-fit metrics measure the ability of an ACI, fitted on a subset of the participant’s data, to predict unseen data from the same participant. Complementary to these “within-participant” cross-prediction measures, it can be useful to assess the goodness of fit of an ACI on a test set extracted from a different participant (“between-participant” cross-prediction). For this purpose, we extended the cross-validation algorithm to allow for the computation of the prediction based on a different ACI. The cross validation can be computed from another participant, as used by Carranante et al. (2024), or from the same participant in a different condition, as used by Osses and Varnet (2024). The cross validations are controlled by the option ‘ACI_crosspred’ indicating the path to the *ACI_*.mat* file to be used for the prediction. The between-participant cross-validation uses the same subsets as the within-participant cross-validation, making it possible to directly compare the estimated goodness-of fit metrics.

Unlike previous validation analyses, the results of the between-participant cross-validation are not saved in the *ACI_*.mat* file. This is because the ACI of the participant is usually computed before running any between-participant analyses and, more importantly, there might be too many ways of cross validate data, so that there is no straightforward way of defining what is the most useful way to provide a cross validation framework that suits the needs of every curent and future user of the fastACI toolbox. Our current solution is to generate a separate file containing the cross-validation data, a *Crosspred_*.mat* file, that will be located (if requested) within the corresponding participant’s *Result* folder.

## 6 CASE STUDIES

In this section, we illustrate the possibilities offered by the toolbox through a series of case studies. These examples demonstrate how the toolbox can be used to replicate existing studies, reproduce published results, and compare different experimental setups, different noise types and different estimation methods. Together, these case studies highlight the toolbox’s versatility and reliability in various research contexts.

### 6.1 Replication of Ahumada et al. (1975)

To illustrate the practical application of the toolbox in the case of simple nonlinguistic stimuli, we describe here the replication of the seminal experiment detailed in Ahumada et al. (1975), considered as one of the earliest examples of auditory reverse correlation studies. At the time, extensive investigation of the perceptual cues underlying tone-in-noise detection had been conducted using conventional psychoacoustic procedures (see in particular Green and Swets, 1966; Sherwin et al., 1956). In 1975, Ahumada et al. addressed this question following a revcorr approach. They analyzed the relationship between fine acoustic details of the noise and subject responses, on a trial-by-trial basis. Doing so, they expected to determine the sound characteristics that are extracted and used by the listener to detect the target. In this section, we re-implement their experiment within the fastACI toolbox to demonstrate the versatility of the framework. We also present the data collected on four participants and compare the results with Ahumada et al.’s original findings.

The experiment is available in the toolbox under the name `replication_ahumada1975`. Similar to the original study, 3200 stimuli were presented consisting in 500-msec Gaussian white noise generated with a sampling frequency of 10 kHz. Half of these stimuli also included a 100-msec, 500-Hz tone, added to the temporal middle of the masker with a fixed signal-to-noise ratio (10 log_10_(*E*_*s*_*/N*_0_) = 11.8 dB). Tone-present and tone-absent stimuli were presented in a random order, through headphones, at a 65 dB sound pressure level. Given the difficulty of the experiment, probe trials with a more favorable SNR (31 dB higher, although note that this parameter was not specified in the original article) were presented every tenth stimulus.

In the interest of the demonstration, we chose to deviate from the original experiment in three respects. First, participants were instructed to provide a yes/no response (tone present or tone absent) instead of a 4-point Likert scale judgment. The use of binary responses was preferred because it aligns with modern reverse correlation studies and it is more compatible with the different post-processing pipelines described in Section 5.3. Second, the initial study included only a single block of 400 stimuli repeated eight times in random order. As this methodological choice was likely guided by computational constraints at the time, we decided to generate 3200 independent stimuli instead, divided into 8 blocks of 400 trials. Finally, stimuli were presented at a level of 65 dB instead of 85 dB, as the latter was deemed too loud by the participants.

As indicated in Section 3.3, the experimental design is entirely described by a set of four scripts (`replication_ahumada1975 cfg.m`, `replication_ahumada1975_set.m`, `replication_ahumada1975_init.m`, `replication_ahumada1975_user.m`), located within the *Experiments* folder of the toolbox. The set-up file contains the generic parameters related to calibration, sampling rate, number of trials and number of targets (this tone-detection task has two targets: tone present and tone absent). As this script is executed only once, it is also used at the same time to generate and store the 500-Hz target tone. The initialization file generates 3200 white noise backgrounds which are stored in a *NoiseStims* folder. The configuration file specifies all experimental parameters, including the number of trials per experimental session, the absence of a warmup phase, the SNR, the fixation of this experimental variable over the course of the experiment (cfg_inout.adapt = 0) except for the easy ‘probes’ every tenth trial (cfg_inout.probe_periodicity = 10). Finally, the user file creates the stimulus for a given trial, depending on the identifier of the corresponding noise background waveform, the SNR, and whether the target is present or not.

In line with the original study, the time-frequency representation of the noise was obtained by computing the energy values in a 5-by-5 matrix defining 25 regions (or “pixels”) with a frequency and time resolution of 50-Hz and 100-ms, respectively. In this representation, the central pixel corresponds to the location of the tone in target-present trials, while the 24 other pixels contain only noise. An ideal observer should therefore pay attention only to the energy in the central pixel. These coarse spectrograms were analyzed together with the behavioral responses through the ‘classical _revcorr’ estimation procedure (Section 5.3). This is mathematically equivalent (up to a scaling factor) to the linear model used by Ahumada et al. This processing of the sounds is carried by an experiment-specific function `replication_ahumada1975_dataload.m` which shadows the default `fastACI_getACI_dataload.m` function.

As the deviations from the original protocol were only minor, we expected to replicate the main result of the original study. Thus, the presence of noise energy in the region of the target tone should correlate positively with the perception of a tone. A similar positive correlation should be found in the segments following the tone, in the same frequency band. Conversely, the correlation should be negative in higher and lower frequency bands during the signal interval.

The resulting images for *N* = 4 participants are shown in Figure 5. The pattern of correlations supports Ahumada et al’s observation that the presence of noise energy on the signal interval (central pixel), but also in the following intervals, biases the listener towards perceiving a tone. Our participants also appear to anticipate the tone to some extent, as the preceding interval at the target frequency is also associated with a positive correlation, a feature that was not present in Ahumada et al’s data. Furthermore, the listeners gave overall negative weight to energy surrounding the target frequency, consistent with the original study. A more detailed description of these replication results is provided in Le Bagousse and Varnet (2025).

**Figure 5.**
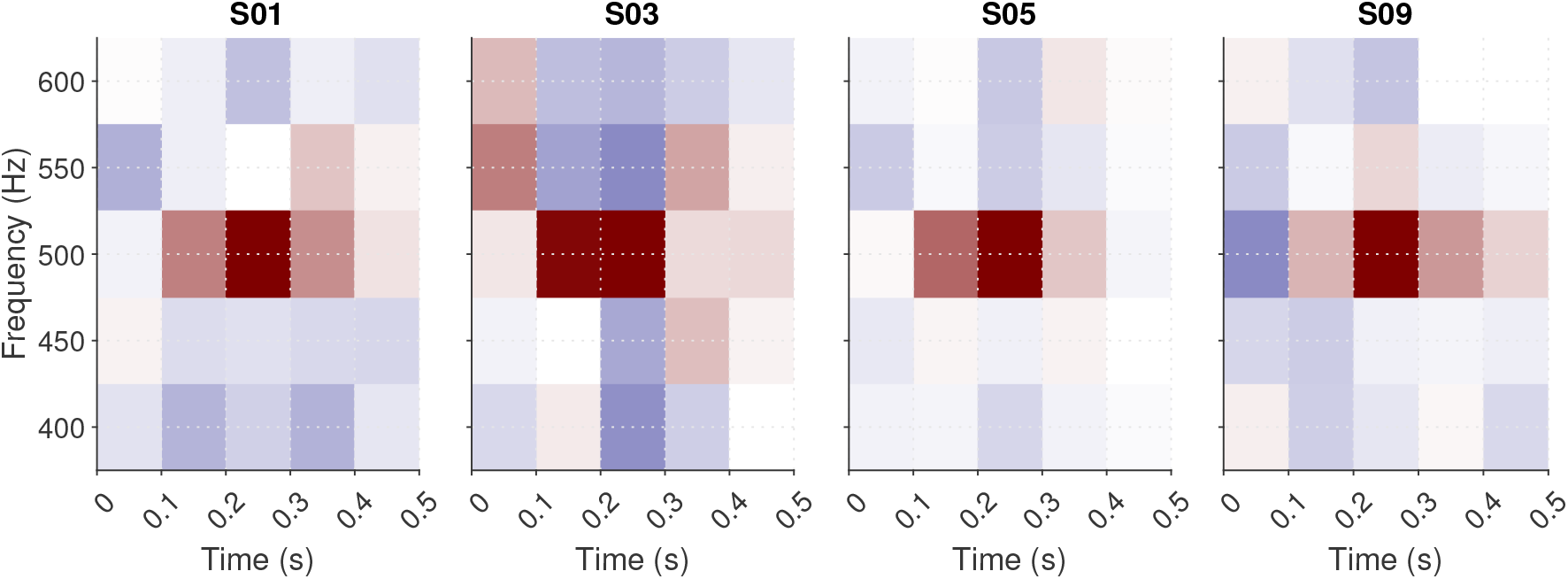
ACIs for four participants in the `replication_ahumada1975` experiment. Red (positive) weights mark regions where the presence of noise energy increases “tone present” responses. Blue (negative) weights mark regions favoring “tone absent” responses. The central pixel indicates the target tone location.

### 6.2 Reproducing published results: Osses & Varnet (2024)

One fundamental goal of a scientific workflow is to ensure that the final publish work is computationally reproducible, that is, that anyone can use the same data to reproduce the same results and figures. The use of an open-source toolbox allows researchers to systematically document their workflows, standardize their procedures, and ensure that all the described analyses can be accurately replicated by others. Using a consistent and transparent set of tools also helps minimizing variability and errors, making it easier to verify findings and build on previous work. A central objectives of the fastACI toolbox is to make it easier to retrieve the experimental sound stimuli, reproduce analyses and re-generate figures. This section provides an example based on one of our recent publication (Osses and Varnet, 2024).

#### 6.2.1 Retrieving the experimental sound stimuli

The first step for reproducing the analysis from a published study is to retrieve the data, in particular the stimuli used in the experiment. In auditory revcorr experiment, this typically involves downloading a large number of .wav files, which can be time-consuming and requires a fast and stable internet connection. The toolbox offers a way to bypass this issue by instead downloading the random seeds, stored in the *cfgcrea*_*\*.mat* file, and recreating the waveforms locally, as decribed in Section 4.4.

The thirty-six datasets used in Osses and Varnet (2024)’s experiments are openly available on Zenodo (https://zenodo.org/records/7476407). To regenerate the sounds using the toolbox, users only need to download the configuration files (available in folder *02-Raw-data.zip*). Then the function `fastACI_experiment_init from_cfg_crea.m` can be executed with the cfg_crea variable as input to regenerate the set of 4000 stimuli corresponding to one participant in one condition. For this particular study, we implemented a script `publ_osses2022b_preregistration_0_init_participants.m` to iterate the procedure on the 12 participants and the 3 conditions.^3^ Once generated, the noises are stored in separate folders within the participant directory named *NoiseStim-white, NoiseStim-bump*, and *NoiseStim-MPS*, each of them containing 4000 waveforms.

The folder *02-Raw-data.zip* also contains the behavioral data collected from each participant, stored in the three *savegame*_*\*.mat* files, corresponding to the three conditions.

#### 6.2.2 Recreating all study figures

Once the data is on disk, all figures from the main text and supplementary materials can be reproduced using the fastACI script `publ_osses2023c_JASA_figs.m`. For instance, Figure 1 from Osses and Varnet (2024) can be obtained with the command `publ_osses2023c_JASA_figs(`‘`fig1`’).

Implementing an analysis workflow as a unique script that can easily be executed by any reader necessarily requires extra effort from the authors. However, the consistent structure of the toolbox ensures that scripts follow a common format, making it straightforward to adapt existing scripts for new analyses.

### 6.3 Effect of target utterances

A limitation of the ACI paradigm, particularly in its application to speech perception, is that each response category (e.g., each phoneme) is typically associated with a limited number of target sounds (e.g., one single utterance of that phoneme). Although it is in theory possible to use multiple utterances of each phonemes, as in Varnet et al. (2015a), this necessarily leads to less clearly defined images when data are aggregated over the entire experiment, as the exact spectrotemporal positions of the cues vary across utterances. Consequently, experimenters usually restrict the number of speech targets to one exemplar of each categories (e.g., one recording of /aba/ and one recording of /ada/, Varnet et al., 2013; Osses and Varnet, 2024). Such a ‘frozen speech’ task is rather unnatural as speech production typically involves a large amount of variability. Furthermore the time-intensive nature of the revcorr paradigm may encourage participants to rely on fortuitous acoustic differences between the speech utterances, rather than on speech cues used in natural listening conditions. It is therefore legitimate to ask how dependent the findings are from the particular acoustic characteristics of the target sounds.

Figure 6 presents the results of the same participant on a fixed phonetic contrast ([aba] / [ada]) using different pairs of utterances. The pairs of targets in ABDA21 and ABDA24 corresponded to natural recordings of /aba/ and /ada/ from OLLO (two different locutors, male and female). The targets in each pair were equalized in syllable duration and intensity. The target pair in ABDA13 was artificially modified: the the initial /a/, produced in isolation, was combined with a recording of /ba/ or a recording of /da/. Therefore, unlike natural speech sounds, the initial vowel in ABDA13 contains no coarticulatory information about the following consonant. The parameters that differ between the three experiments are indicated in Table 4. The time-frequency representations of the targets used in each of the three experiments are presented in Figure 6 together with the resulting ACIs estimated using a GLM with L1 regularization on a Gaussian basis.

**Table 4.** List of parameters used in each [aba]-[ada] discrimination experiment considered. The adaptive procedures used in different experiments targeted the same overall performance score of 70.7%. Further details are given in the body text. *N*_*trials*_ = total number of collected trials.

| Exp. name | Masker type | Speaker | $N_{trials}$ | Staircase rule | Roving | Ref. |
| --- | --- | --- | --- | --- | --- | --- |
| ABDA24 | white noise | male 1 | 4000 | weighted 1-up 1-down | $\pm 2.5$ dB | Osses and Varnet (2024) |
| ABDA21 | bump noise | female 2 | 5000 | weighted 1-up 1-down | no | - |
| ABDA13 | white noise | female 1 | 10.000 | transformed 1-up 2-down | no | Varnet et al. (2013) |

**Figure 6.**
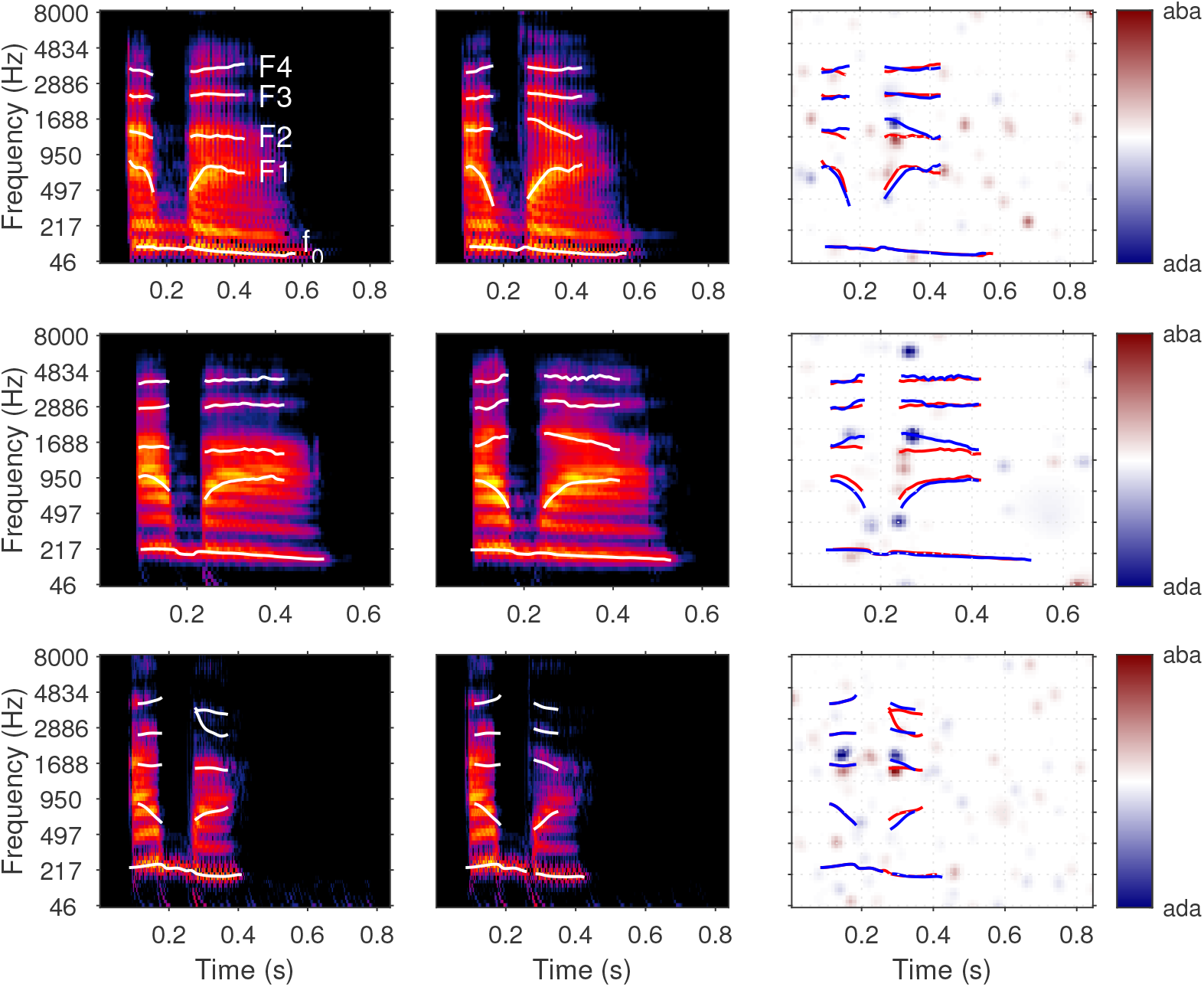
Results of three ACI experiments performed on different [aba]-[ada] pairs by the same participant. The rows correspond to the three experiments described in Table 4, with the time-frequency representations of the targets in the first ([aba]) and second ([ada]) columns, and the ACI obtained in the third column. ACIs are calculated using a penalized GLM with L1 regularization on a Gaussian basis, and normalized in maximum absolute value. Formant and *f*_0_ trajectories are indicated on the spectrograms, and they are reproduced on the corresponding ACI to facilitate interpretation.

The ACIs for experiments ADBA13, ABDA24, and ABDA21 reveal a clear pattern of weights, confirming that the participant is actively extracting information from the noisy stimuli to perform the task. For each of the three experiments, the strongest weights are localized in the time-frequency regions corresponding to the second formant (*F*_2_) transition, at the onset of the second syllable. More precisely, all ACIs showed a similar pattern of weights in this region, organized vertically, with a positive (red) cluster below a negative (blue) cluster. This results, consistent with the obtained by other participants in the same task (Varnet et al., 2013; Osses and Varnet, 2024; Carranante et al., 2024), confirms the critical role of the *F*_2_ onset as a cue for [b]-[d] categorisation, as demonstrated using other psycholinguistic methods (Liberman et al., 1952). Therefore, despite the natural variability between speech targets (e.g., variability in the exact spectrotemporal positions of the *F*_2_ onset), the method is able to reliably identify the primary speech cue for the considered phonetic contrast.

Although the *F*_2_ onset cue is identified in all three experiments, the ACIs also reveal some variability in the listening strategies used by the same participant when performing the same task with different /aba/-/ada/ pairs. The listener seems to be able to rely more or less on secondary speech cues to distinguish between targets, including the *F*_1_ transition near the onset of the second syllable (ABDA24 and ABDA21), the presence of high frequency energy within the intervocalic interval (ABDA21), matching the spectrotemporal position of a potential consonant release burst, and the *F*_2_ transition at the offset of the first syllable (ABDA13). The perceptual weights associated to these cues, when present, are weaker and therefore they appear as a secondary source of information for performing the tasks (see Carranante et al., 2024, for a discussion of these cues). Listeners are able to adapt their weighting strategy depending on the availability and robustness of a cue. It is therefore no surprise that the auditory revcorr method applied to different pairs of targets results in different sets of weights. For example, the use of a coarticulation cue in the first syllable may depend on the relative energy (and therefore the audibility) of this syllable, low in ABDA2023 but high in ABDA2013 (Osses et al., 2022a).

The main cue on *F*_2_ seems to be present in all three experiments, while only secondary cue weights are subject to variability.^4^ This leads to the question whether the strategies revealed with the ACI method are entirely contingent on the choice of particular targets, as the listener would be able to learn target-specific cues over the course of the experiment. In particular, do they depend on the presence/absence of a particular cue in the target? The case of experiment ABDA13 is particularly interesting in that respect. As noted above, in this experiment the initial vowel contains no relevant information for the task, unlike natural speech sounds, as both targets begin with exactly the same /a/. Nevertheless, the ACI shows large weights in the *F*_2_ region in this segment. This highlights an important property of the revcorr method: it can reveal cues that listeners expect to find in the stimuli, even if they are not actually present in the chosen set of targets. This property has already been demonstrated in visual psychophysics (Gold et al., 2000; Gosselin and Schyns, 2002, 2003). In particular, the so-called “superstitious” revcorr protocols in which the stimuli contain only noise (without any target) nevertheless allow for the calculation of stable kernels, reflecting the expectations that participants “project” onto the stimuli (Gosselin and Schyns, 2003; Liu et al., 2014). In other words, the results of the ABDA2013 experiment illustrates that participants are not able to adapt, over the course of the experiment, to the absence of a particular cue in the target.

In conclusion, this short case study illustrates that, while the ACIs depend on the particular choice of targets, the listeners’ ability to adapt their strategies to extract additional, superficial, acoustic cues seems to be limited and does not directly challenge the efficacy of the method. The ACI method can reveal the acoustic cues that are audible, or expected to be audible, in the targets, but it does not necessarily provide an exhaustive list of all possible cues used for a given contrast.

### 6.4 Comparing between different noise types

In this section, we focus on the effect of the statistics of the background noises with respect to the efficiency and robustness of the method. Three different noise types are considered that have a flat long-term spectrum, but differ in the amount of temporal envelope fluctuations: White noise (WN), white noise low-pass filtered in the modulation-power-spectrum (MPS) domain, and bump noise (BP) (see Osses and Varnet, 2024, for a description of these three types of masker). In Osses and Varnet (2024) *N* = 12 participants performed 4000 trials of the same [aba]-[ada] categorization task in each of the three noise conditions (12000 trials / participant). The averaged ACI obtained in each noise condition are shown in Figure 7.^5^ The out-of-sample prediction accuracy of the GLM fitted using the ‘glm_L1_GB’ option was measured separately for the three noise conditions (see Section 5.4.3). The findings indicated that the GLM reached higher prediction performances in the BP and MPS noise conditions compared to the WN condition (13.3% and 11.8% vs. 8.1% above chance level, respectively). This suggests that noise maskers with larger modulations in the low-frequency range (BP and MPS) interfere with the phoneme categorization process in a more systematic way (i.e., yielding more predictable errors), therefore potentially resulting in a more robust estimation of the ACI.

**Figure 7.**
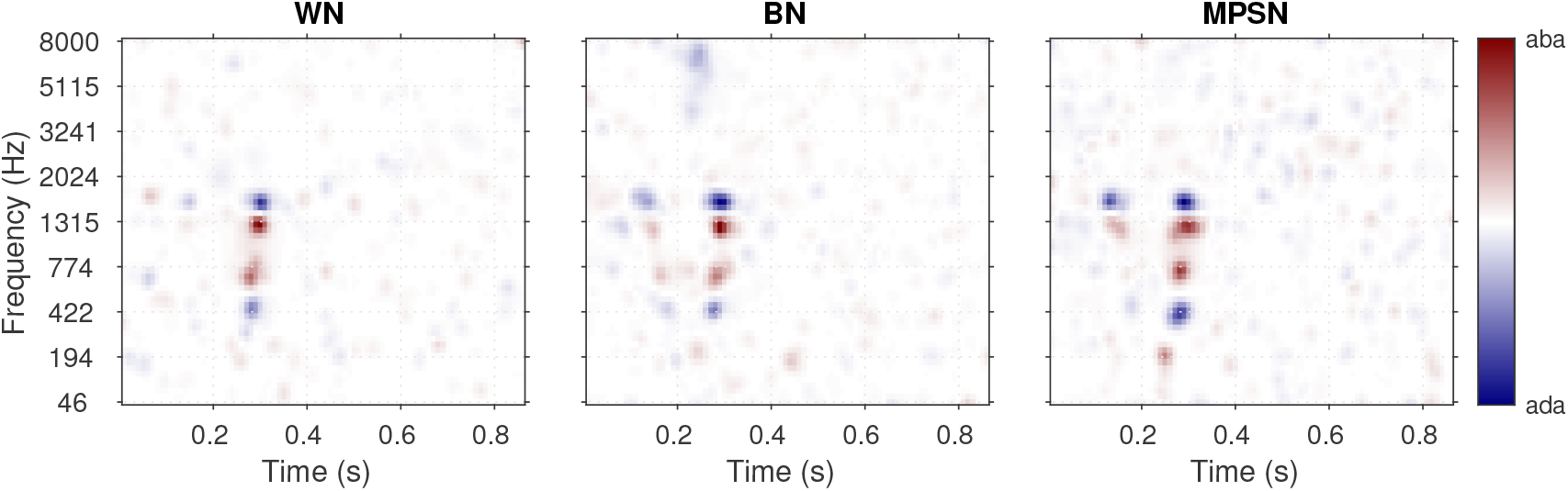
Mean ACI across 12 participants, obtained for each of the three noise conditions : white noise (WN), bump noise (BN), MPS noise (MPSN).

In this case study, we confirm this finding using two additional metrics. For this purpose, we re-analyzed the data from Osses and Varnet (2024). In order to quantify the speed of convergence, each dataset (one participant in one condition) was first randomized, then divided into 10 subsets corresponding to the first *k* blocks of 400 trials (total number of trials *n* = *k ·* 400 with *k ∈* [[1, 10]]). For each subset, a “partial” ACI was derived using the ‘correlation’ approach – see the following section for the results of the ‘glm_L2’ and ‘glm_L1_GB’ options. For each of these partial ACIs, two metrics were computed: the correlation with the final ACI and the cue-to-noise ratio.

1. **Correlation with the final ACI**: In order to quantify the speed of convergence, the similarity of each partial ACI with the final ACI (corresponding to *n* = 4000) was measured using Pearson correlation. This is a widely used and straightforward metrics for assessing the convergence of the revcorr method (e.g. Burred et al., 2019; Varnet et al., 2013). However a major downside of this metrics is that it assumes that the final ACI correspond to the “true” template, which is rarely the case in practice. As a result, all correlation metrics necessarily converge to 1, making it difficult to compare the accuracy of convergence between different conditions.
2. **Cue-to-noise ratio**: Although a direct comparison between the estimated template and the true template is usually not feasible, as the true template is generally unknown, in this specific task we can capitalize on our understanding of relevant cues to assess the accuracy of the estimation. Specifically, in [b]-[d] phoneme discrimination tasks, the *F*_2_ onset is known to play a major role (see previous section). This finding has been replicated many times in the psycholinguistic literature, and the F2 cue was observed in every participant in our previous ACI studies (Osses and Varnet, 2024; Carranante et al., 2024; Varnet et al., 2013). Conversely, no cue is expected to be found in the same frequency range during the silent segment after the end of the second syllable. Therefore, we define the cue-to-noise ratio as the mean squared weights in the time-frequency region of the *F*_2_ onset (1 to 2 kHz, 0.25 to 0.3 s) divided by the mean squared weights in the non-relevant time-frequency region (1 to 2 kHz, 0.55 to 0.8 s).

The comparison of the three different noise types in terms of correlation analysis and cue-to-noise ratio yielded results consistent with the prediction accuracy metrics (Figure 8). The reverse correlation method applied to a consonant-in-noise discrimination task converges more quickly and more robustly to a stable template when the background noises contain dominant components in the low modulation frequency range (between 0 and 40 Hz), which is the case for MPS and BP noises. The prominent envelope fluctuations in this range lead to more systematic confusion errors compared to white noise and, therefore, to higher prediction accuracy and more robust reverse correlation results. In particular, the BP noise yields a slightly better cue to noise ratio (= 3.2 at *n* = 4.000 trials) compared to the two other types of noise (2.4 for MPS, 1.8 for WN). The same conclusion was reached using other fitting procedures (see following section).

**Figure 8.**
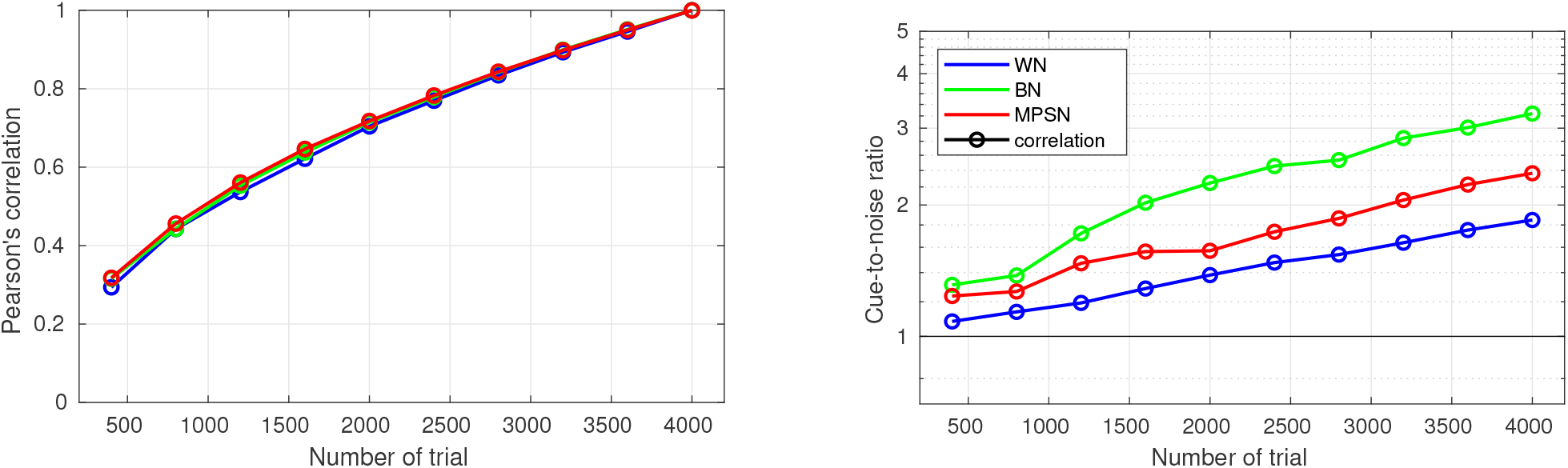
Convergence of the estimated ACI throughout the experiment, for the three types of noise. All ACI are computed on the same group of 12 participants from Osses and Varnet (2024), using the ‘correlation’ approach. Left: Correlation between the “partial” ACI derived using *n* trials and the ACI derived using all trials (*n* = 4000). Right: Cue-to-noise ratio for each partial ACI. The black line corresponds to a cue-to-noise ratio of 1, indicating that relevant and non-relevant time-frequency regions of the ACI are associated with the same absolute amplitude of weights.

### 6.5 Comparing between different estimation methods

In this section, the same set of data used in paragraph 6.4 was subjected to three separate pipelines of analysis. All analysis parameters were the same except for the fitting method which was set to one of the three options presented in Section 5.3: ‘correlation’, ‘glm_L1_GB’ or ‘glm_L2’.

Two main conclusions can be drawn from Figure 9. First, for small datasets (here approximately *n* < 2400), only the standard correlation procedure is able to produce a reliable result. The obtained templates are only slightly to moderately correlated with the final ACI, with a low cue-to-noise ratio in the range 1-3. This approach can still be useful for experiments that are interested in gathering data from a large sample of participants, even at the expense of the quality of individual data. In this case, it is still possible to post-process the estimated images to improve the quality (for example using a simpler two-dimensional smoothing).

**Figure 9.**
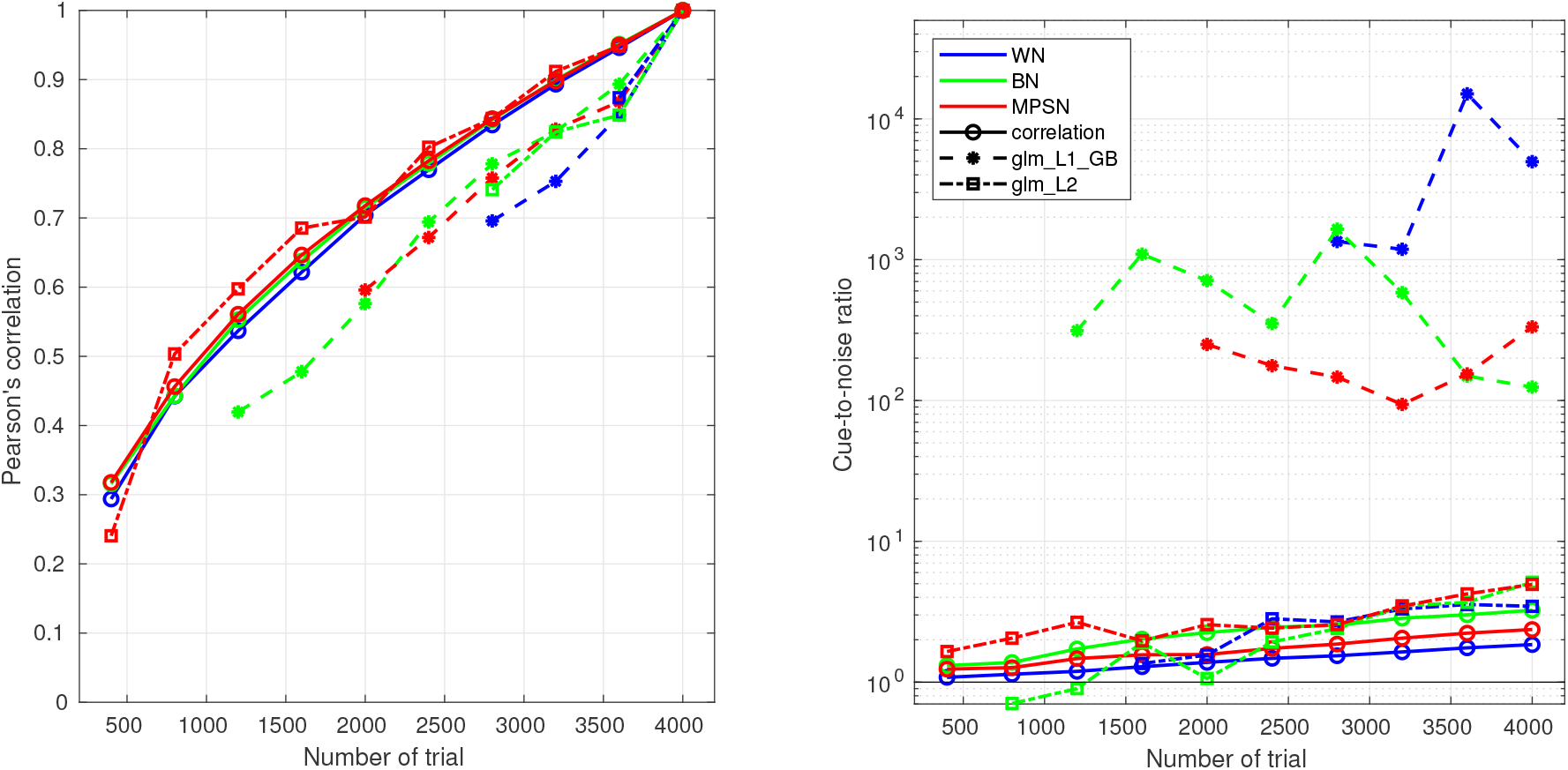
Convergence of the fit throughout the experiment, for the three estimation methods. Same legend as in Figure 8. For penalized regression methods (glm_L2 and glm_L1_GB), only datapoints corresponding to estimations that successfully converged for at least 8 of the 12 participants are shown.

On the contrary, the ‘glm_L1_GB’ and ‘glm_L2’ approaches require a minimum of *≈* 2400 trials to ensure the convergence of the hyperparameter. When this condition is met, they consistently produce better estimates that the `correlation` approach, with an improvement of the cue-to-noise ratio by a factor 2 in the case of ‘glm_L2’, and by a factor of 12 or higher in the case of ‘glm_L1_GB’. While this finding may appear to support a preference for using the option ‘glm_L1_GB’ over ‘glm_L2’, it is crucial to acknowledge that this conclusion is contingent upon the adequacy of the prior for the specific task and targets considered. In the present case, as highlighted in Section 5.3, the ‘glm_L1_GB’ is more appropriate as the acoustic cues for stop consonant discrimination are highly localized in the time-frequency space.

## 7 CONCLUSION

The fastACI toolbox implements a type of psychoacoustics experiments that have been used in auditory science for over fifty years. During this time, revcorr experiments have often been conducted with a wide range of parameters, both in terms of experimental design (e.g., yes/no or two-intervals forced-choice tasks, psychophysical staircases or constant stimuli, background noise or random signal modifications) and analysis methods (e.g., spectrogram or auditory-based representation, simple linear regression or regularized GLM). This lack of standardization has made it difficult to compare results across studies and has limited reproducibility. By integrating these options within a consistent and flexible framework, fastACI enables, for the first time, systematic comparisons across different configurations. The toolbox is also designed to be as “plug-and-play” as possible, allowing researchers unfamiliar with the reverse correlation approach to easily apply it to their own research questions. Finally, fastACI is open source, supporting transparent, replicable, and computationally reproducible research.

## AUTHOR CONTRIBUTIONS

L.V. and A.O implemented the toolbox and the analysis scripts and wrote the original draft of the manuscript. L.V., A.O., and A.L.B. collected the data in the different experiments, and revised the manuscript.

## DATA AVAILABILITY STATEMENT

The datasets analyzed for this study can be found on Zotero in the following repositories: https://zenodo.org/records/14972392 for section 6.1, https://zenodo.org/records/7476407 for sections 6.2, 6.3, 6.4, and 6.5.

## REPLICATING THE FIGURES AND ANALYSES

After downloading the data (see above), all figures and analyses can be replicated using the fastACI toolbox (Osses and Varnet, 2021b) as of version 1.4. Figures 3 to 9 from the main text can be obtained by running the script `publ osses2025 bioRxiv figs.m`, indicating the corresponding figure number.

## Footnotes

1 Strictly speaking, however, the auditory revcorr approach is not limited to the auditory classification image but also encompasses other methods such as that of auditory bubbles (Mandel et al., 2016; Venezia et al., 2016).

2 Although multicollinearity is often treated as a challenge for accurate estimation, it is, in fact, fundamental to the principle of statistical control.

3 Note that for recreating the MPS noises, the PhaseRet toolbox needs to be installed and compiled. No extra dependencies are required to reproduce the white and bump noises.

4 The same is true with respect to interindividual variability, as noted in Carranante et al. (2024).

5 Although the auditory system is generally able to adapt its cue-weighting strategy to the statistics of the background noise, assigning higher weights to more robust cues, it is noteworthy that the three ACIs are quite similar to each other. This similarity is likely due to the fact that all three masker types have a white-noise-like long-term spectrum, resulting in similar local SNR for each cue across the three conditions. The only exception is the bump noise, which shows a larger weight in the burst cue region for the bump noise. This may be because the bumps in the masker are more likely to be confused with a release burst sound.

